# PIEZO1-mediated mechanosensation links aging to bladder dysfunction

**DOI:** 10.64898/2026.05.24.726025

**Authors:** Yasmeen M F Hamed, Vikram Joshi, Kevin Wilhelm, Luis O Romero, Olivia D Solomon, Tammy B Kwok, Eskarleth D Lopez Gonzalez, Monica M Ridlon, Shriya Pendyala, Catherine M Calhoun, Alexa E Martinez, Jennifer K Asmussen, Kimberly Keil Stietz, Chad M Vezina, Valeria Vásquez, Olivier Lichtarge, Arthur Beyder, Kara L Marshall

## Abstract

Aging is accompanied by profound changes in bladder function, leading to increased urinary frequency and incontinence that impair quality of life in humans and are recapitulated in mouse models. Bladder filling and emptying rely on precise mechanosensory feedback, yet how aging alters this sensory control remains unclear. PIEZO ion channels convert mechanical forces into cellular signals essential for bladder fullness sensation. Using inducible smooth-muscle specific *Piezo1* deletion, we find that loss of *Piezo1* attenuates aging-related bladder dysfunction. Dietary enrichment with margaric acid, a membrane-active fatty acid previously shown to inhibit PIEZO channels, reduced urinary dysfunction in aged mice. In humans, genotype–phenotype analyses reveal an association between a *PIEZO1* gain-of-function variant and early-onset neuromuscular bladder dysfunction. Together, these findings define a smooth-muscle, PIEZO1-mediated mechanosensory basis for aging-related bladder dysfunction and introduce a non-invasive strategy for targeting PIEZO channels *in vivo*.

## INTRODUCTION

Mechanosensation is the ability of cells and tissues to detect and respond to mechanical stimuli—a process that is fundamental for normal function of many organ systems^1^. From touch sensation and proprioception to vascular tone, gastrointestinal motility, and urinary function, mechanically evoked signaling coordinates physiological responses essential for survival^1^. Many of these processes decline with age, contributing to impaired mobility, sensory dysfunction, and visceral organ dysregulation^2^. In the lower urinary tract, aging is associated with increased frequency, urgency, and incontinence, significantly diminishing quality of life and impeding independence^3^. Although these symptoms are well described in humans and recapitulated in aged mice^4,5^, the tissue-level mechanisms that drive aging-associated bladder dysfunction remain poorly defined. Specifically, it is unknown whether and how altered mechanotransduction within detrusor smooth muscle directly contributes to defective bladder storage and voiding. Multiple bladder cell types are mechanosensitive: urothelial cells, smooth muscle cells (SMCs), and innervating sensory neurons all contribute to the stretch responses essential for bladder function^6–8^. PIEZO ion channels are key mechanosensors that convert mechanical stimuli into cellular signals essential for numerous physiological processes^1,9,10^. Previous work has shown that PIEZO2 is critical for bladder fullness sensation and urinary reflexes, and both PIEZO1 and PIEZO2 contribute to urothelial cell signaling, which communicates with the underlying muscle cells and sensory neurons^6,7,11,12^. PIEZO1 is more broadly expressed in bladder tissues, including in SMCs^13^. Aging-related changes in neuronal bladder reflex function and urothelial signaling during filling have been implicated in aging-related urinary deficits^14–17^. While some aging-associated structural changes in the bladder smooth muscle itself have also been previously described^18,19^, mechanosensory changes are not well characterized. Both bladder functional and sensory properties substantively contribute to mediating proper continence and voiding: its ability to expand and accommodate urine without a significant rise in pressure allows for storage without prematurely inducing a sense of urgency, and the effectiveness of contraction controls efficient urinary output. PIEZO1’s role in the bladder detrusor smooth muscle is unknown, but it is an important candidate for mediating smooth muscle mechanosensory processes. We hypothesized that PIEZO1 activity in the detrusor smooth muscle might underlie aging-associated declines in bladder function.

Here, using genetic mouse models, we uncover a specific requirement for PIEZO1-mediated smooth muscle mechanosensation in aging-related bladder dysfunction. We also identify a targeted dietary intervention enriched with the saturated fatty acid, margaric acid, as a potent modulator of mechanosensory signaling *in vivo*, corroborating previous *in vitro* findings^20,21^. We show that margaric acid inhibits mechanically gated currents in dorsal root ganglia (DRG) sensory neurons from young mice fed the margaric acid-enriched diet and markedly improves urinary function in aged mice. This intervention reduces voiding frequency and enhances continence, with physiological evidence supporting improved detrusor smooth muscle function as a contributing mechanism. Finally, human genotype–phenotype analyses reveal an association between a *PIEZO1* gain-of-function variant that is highly prevalent (9-23%) among individuals of African descent^22^, and earlier-onset neuromuscular bladder dysfunction, suggesting conserved mechanistic principles for the role of mechanosensation in urinary dysfunction across species.

## RESULTS

### Aged mice exhibit dysfunctional urinary behavior

To characterize aging-associated changes in urinary voiding behavior, we performed 4h void spot assays in young adult (8-12 weeks) and aged (92-93 weeks) male mice (Figure 1A). Consistent with prior reports^16^, young mice predominantly voided along the corners and edges of the cage while largely avoiding the center (Figure 1B–C). In contrast, aged mice exhibited increased voiding frequency and a higher number of voiding events in the cage center (Figure 1C), despite having comparable total urination (Figure 1D), a pattern consistent with impaired urinary control or an incontinence-like phenotype^16^. The increase in voiding frequency (Figure 1E) was driven primarily by an increase in smaller (secondary) voids (Figure 1F–G). These differences were not attributed to altered exploratory behavior, as young and aged mice spent an equivalent fraction of time in the center of the testing arena (Figure 1H–I). These data indicate that aging is associated with altered urinary voiding patterns characterized by increased frequency and incontinence-like patterns in mice, which are also common characteristics of aging-related urinary dysfunction in humans^23^.

**Figure 1.**
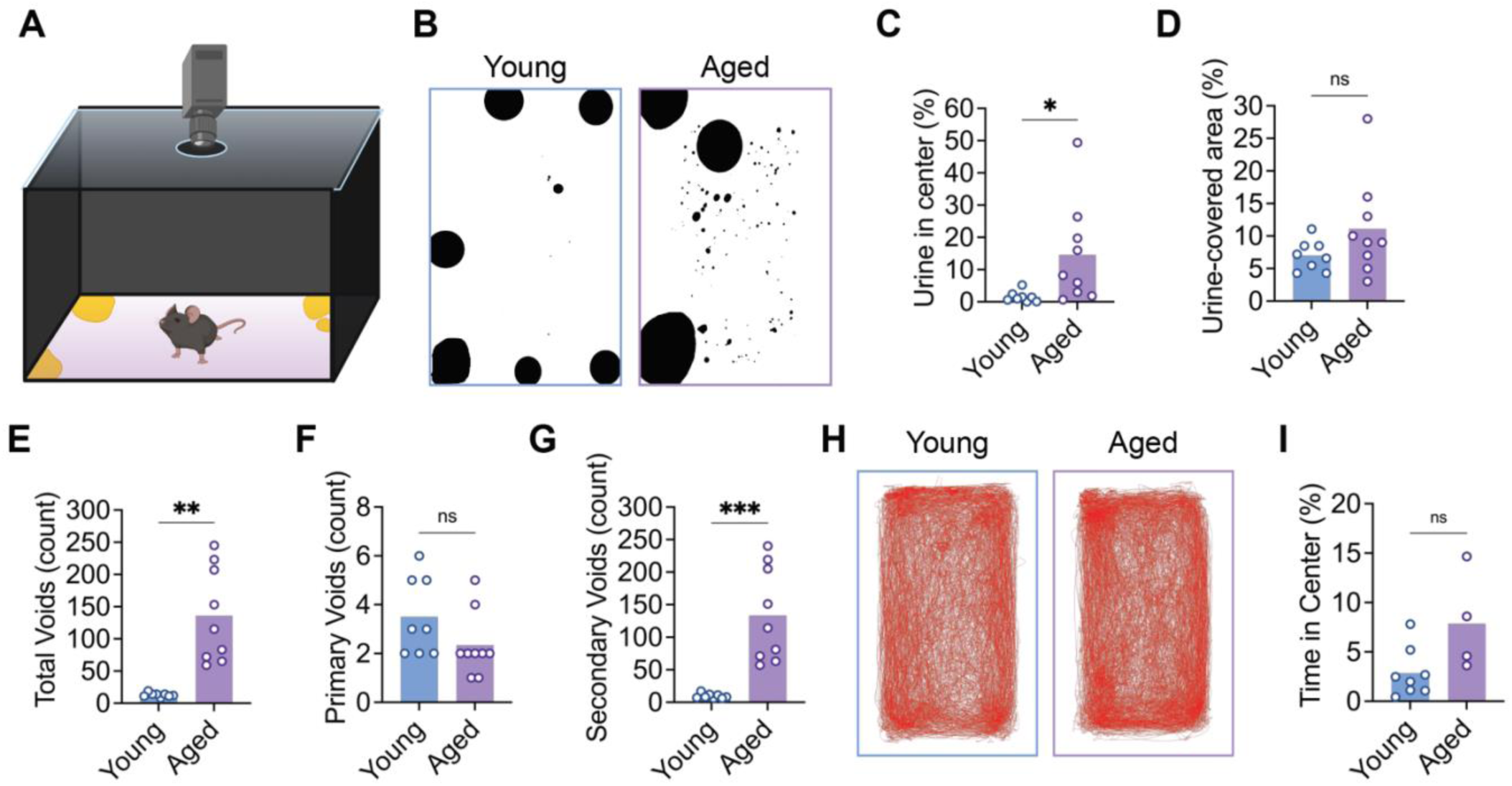
Aged mice exhibit dysfunctional urinary behavior. (A) Illustration showing the void spot assay set-up (4 hours). (B) Representative end-point images of 4 h void spot assay papers from male mice; young (*left*, blue frame) and aged (*right*, purple frame); imaged under UV light. Urine void spots are shown in black; quantified in C-G (n=8-9/group). (C) Percentage of urine deposited in center 25% of cage area in a 4 h void spot assay. (D) Total urine-covered area at the end of a 4 h void spot assay, quantified as percentage of total cage area. (E) Total number of urinary void spots deposited in a 4 h assay, further stratified by size into primary (F) and secondary (G). (H) Line plots showing representative movement traces of young (*left*, blue frame) and aged (*right*, purple frame) mice during the 4 h void spot assay. (I) Percentage of time spent in the center 25% of cage area during the 4 h void spot assay, quantified from movement tracking videos; n=4-8/group. For all bar plots: bars show mean; dots show individual animals. (C-G, I) Statistical test: Welch’s *t*-test. (ns) *p* > 0.05, (*) *p* ≤ 0.05, (**) *p* ≤ 0.01, (***) *p* ≤ 0.001, (****) *p* ≤ 0.0001.

Small, frequent voids in the cage center could indicate loss of volitional control, loss of continence, increased sensations of urgency or an over-active bladder like phenotype in aged males. We did not observe similar changes in *nulligravida* (never pregnant) aged female mice at the same age (92-93 weeks), but only subtle changes starting at 97 weeks of age (Figure S1). Therefore, we focused on males for the rest of the study to characterize the key mechanisms underlying aging-related bladder dysfunction. We first asked whether alterations in innervation are associated with this behavioral change but found no significant difference in overall innervation in young versus aged mice, as observed in both tissue sections (Figure S2A–C) as well as whole-mounts (Figure S2D–E). The mechanosensitivity of neurons and urinary reflexes are ultimately driven by the tissue mechanics at nerve endings embedded within the bladder^24^. Furthermore, previous studies have shown that aging in mouse bladders is associated with impaired detrusor contractility, altered calcium signaling, reduced cholinergic and β–adrenergic receptor expression, increased collagen deposition, and structural remodeling of the muscle wall, which are tied to age-related alterations in voiding behavior^25–28^. Thus, we focused on understanding how the bladder wall muscle itself was changing with age.

### Conditional smooth muscle *Piezo1* deletion protects against aging-related urinary dysfunction and preserves bladder contractility

Smooth muscle cells (SMCs) contribute to aging-related dysfunction of contractile organs, potentially through impaired mechanosensation^29^. Consistent with the idea that mechanosensation plays a critical role in smooth muscle function, recent studies^13^ and our own data (Figure S3A) demonstrate that bladder SMCs express *Piezo1*. To determine whether SMC-expressed *Piezo1* contributes to aging-related bladder dysfunction, we performed void spot assays^30^ in SMC-specific *Piezo1* knockout mice (*Myh11-creERT2^+^; Piezo1^fl/fl^*, induced with Tamoxifen at 7-8 weeks; hereon referred to as P1KO). We confirmed that induced loss of SMC *Piezo1* persists with age and does not impact the expression of *Piezo2* in compensatory mechanisms (Figure S3A). Similar to wildtype animals, both young control (*Piezo1^fl/fl^*) and young P1KO mice primarily voided at the chamber periphery (Figure 2A, B). In contrast, aged control mice exhibited a marked increase in center voiding events, voiding frequency, and small (secondary) voiding events, consistent with incontinence-like phenotype^30^ (Figure 2C, E, G). Despite these changes, aged control mice maintained consistent total urine output and large void spot counts, indicating that primary voiding capacity remains intact (Figure 2D, F). Strikingly, aged P1KO mice, with *Piezo1* deletion induced in young adulthood to allow for normal development, were protected from these deficits and maintained youthful micturition patterns even at 24 months of age (Figure 2C–G). Compared with aged controls, aged P1KO mice showed reduced central voiding while maintaining peripheral voids and normal large-void counts (Figure 2B–C, F). Together, these findings indicate that SMC-PIEZO1 is a key contributor to aging-related urinary dysfunction and demonstrate that inhibition of PIEZO1 could be a viable approach to restore normal bladder function.

**Figure 2.**
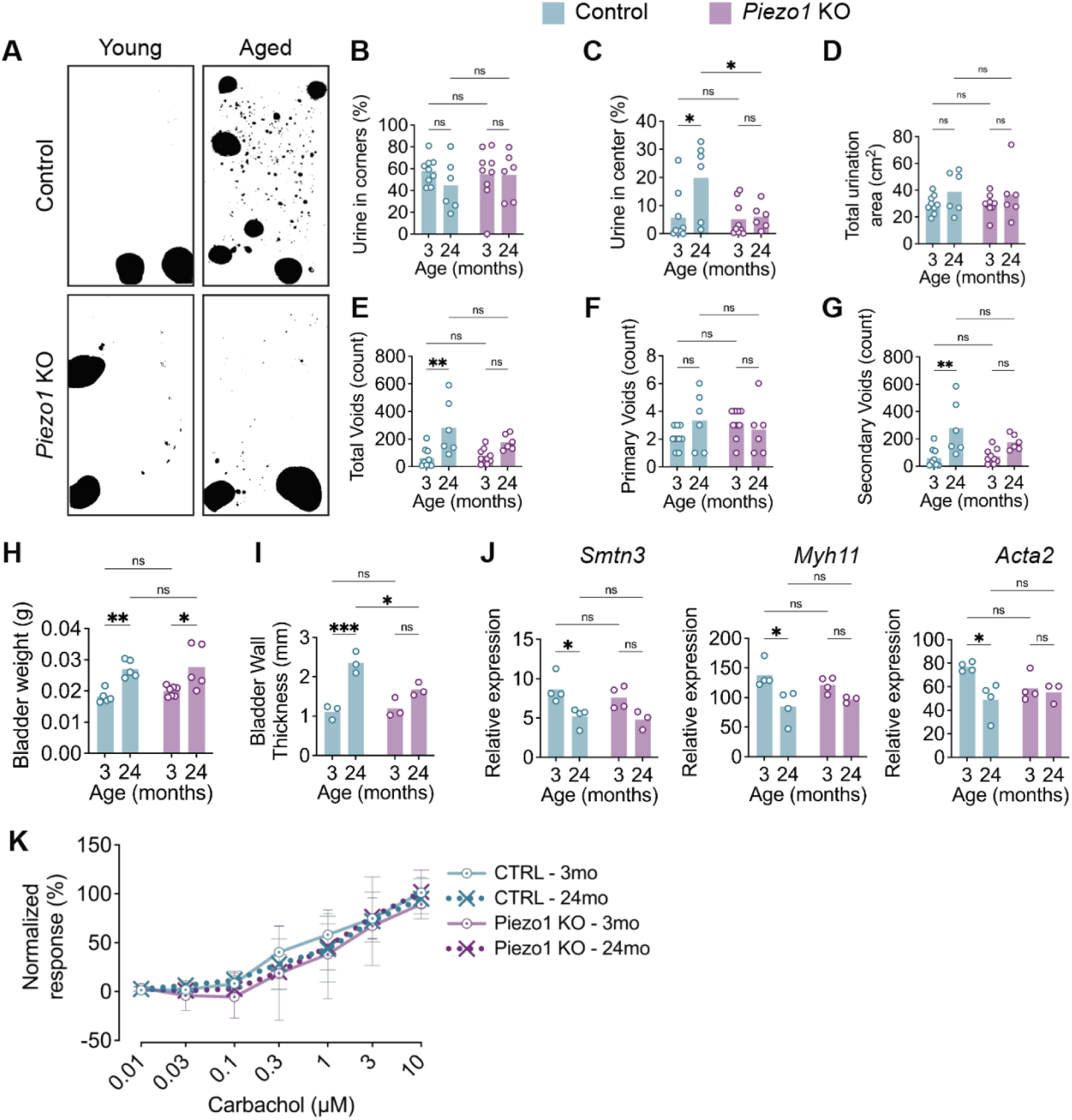
*Piezo1* deletion protects against aging-related urinary dysfunction and preserves bladder contractility. (A) Representative 4h VSA images from control young (*top left*), control aged (*top right*), P1KO young (*bottom left*), and P1KO aged (*bottom right*) mice; black spots = urine. (B-C) Percentage of urine deposited in periphery of the arena (B) and the center 30% of arena (C); n=6-9/group. (D) Total urine-covered area at the end of a 4 h void spot assay, quantified as percentage of total cage area; n=6-9/group. (E) Total number of urinary void spots, further stratified by size into primary (F) and secondary (G). n=6-9/group (H-I) Measurement of bladder physical properties: (H) Wet bladder weights; n=5-7/group and (I) bladder wall thickness; n=3/group. (J) Relative expression of contractility markers (*Smtn3, Myh11, Acta2*) in bladder smooth muscle, normalized to reference genes; n=3-4/group. (K) Bladder contractile response to carbachol, normalized to KCl-induced maximal response. n=4-7/group. For all bar graphs: bars show mean, dots show individual animals; for (K): dots show means ±SD (error bars). Statistical tests: (B-J) Two-way ANOVA with Šídák’s multiple comparisons test. (K) Two-way mixed-effects model (REML) with Tukey’s multiple comparisons test; K, no significance bars = not significant. (ns) *p* > 0.05, (*) *p* ≤ 0.05, (**) *p* ≤ 0.01, (***) *p* ≤ 0.001, (****) *p* ≤ 0.0001.

Aging bladders undergo alterations in structure, biomechanical properties, and urothelial and detrusor function—all of which were shown to affect urinary function in both mice^16,25,27^ and humans^3,31,32^. To define how *Piezo1* deletion alters bladder physiology, we next examined bladder anatomy, physical properties, and contractility using *ex vivo* preparations. Bladder weight increased with age in both control and P1KO mice (Figure 2H). However, age-related bladder wall thickening was observed only in control mice, whereas P1KO bladders remained comparable to their young controls with age and were significantly thinner than aged wildtype controls (Figure 2I), indicating protection from age-related tissue remodeling. At the transcription level, aging decreased the expression of contractile genes (*Smtn3, Myh11, Acta2*) in control bladders, whereas contractile gene expression remained stable in P1KO bladders (Figure 2J), suggesting that PIEZO1 negatively regulates contractile gene expression during aging. To assess contractility in response to cholinergic stimulation, we used carbachol, a stable muscarinic receptor agonist, and observed no change in bladder contractile response across genotypes or with age (Figure 2K). These data are consistent with preserved contractile potential in the absence of PIEZO1 during aging. Aging also did not alter the expression of muscarinic receptors (*Chrm2, Chrm3*) (Figure S3B) or major potassium channels (*Kcnma1, Trek1, Trek2, Traak*) in either genotype (Figure S3C), indicating that PIEZO-mediated mechanosensation does not predominantly drive receptor signaling or SMC excitability. Interestingly, what appears to be altered in aging through PIEZO-dependent signaling is contractile machinery (Figure 2J), which leads to compensatory remodeling (Figure 2I) that in the long-term could underly aging-related pathology.

Direct smooth muscle depolarization via potassium chloride (KCl) and electrical field stimulation (EFS)-evoked contractions were not significantly altered between genotypes or with age (Figure S3D–E), indicating basal muscle strength and neuromuscular transmission remain intact. Similarly, elastic modulus and ATP-induced contractions were not altered by age or genotype (Figure S3F–G). These findings collectively demonstrate that the functional effect of *Piezo1* deletion is primarily preservation of smooth muscle-intrinsic contractile properties in aging rather than overall muscle health or passive mechanical properties. Furthermore, our data indicate that deletion of smooth muscle *Piezo1* prevents the emergence of urinary dysfunction during aging in the bladder, limits structural remodeling, preserves contractile gene expression and intrinsic contractile capacity.

### Dietary margaric acid supplementation inhibits mechanically-gated currents

Mechanical properties of biological tissues are shaped by multiple interacting factors, including cytoskeletal architecture, extracellular matrix interactions, mechanosensitive ion channels, and the lipid composition of the plasma membrane^33–36^. Mechanosensitive PIEZO ion channels are directly gated by membrane tension^37–39^ and are therefore highly sensitive to the biophysical properties of the lipid bilayer^20,21,40,41^. Because pharmacological tools to selectively modulate mechanosensory ion channels *in vivo* are limited, we sought to modulate mechanosensation by altering membrane mechanical properties. As fundamental building blocks of membrane lipids, specific dietary fatty acids are incorporated into cell membranes, where they remodel membrane biophysical properties *in vitro*, *ex vivo*, and *in vivo* and thereby tune the gating of mechanosensitive PIEZO ion channels^20,21,40–42^. Margaric acid, a saturated fatty acid found in dairy products and other foods, increases membrane stiffness and inhibits both PIEZO1 and PIEZO2 channel activity *in vitro* and *ex vivo*^20,21^. We therefore investigated whether a diet enriched in margaric acid could suppress mechanosensory ion channel activity *in vivo* as a potential non-invasive strategy to ameliorate aging-related bladder dysfunction. To assess this in an established control tissue with broad expression of *Piezo2* (and, to a lower extent, *Piezo1*)^43^ that is also amenable to cell membrane manipulation using dietary fatty acids, we utilized dorsal root ganglia (DRGs)^20,21,40,41^. After margaric acid-enriched dietary supplementation (59% of calories from margaric acid), we determined a robust increase in margaric acid content in DRG as well as bladder tissues compared with animals fed standard chow (Figure S4A). These data indicate that dietary supplementation provides a robust and reliable means of enriching peripheral tissues with margaric acid.

To assess the functional consequences of margaric acid enrichment on mechanosensory signaling, we performed whole-cell patch-clamp electrophysiology in DRG sensory neurons from mice fed an isocaloric control diet (59% calories from anhydrous milk fat), mice fed the MA diet, and mice fed the control diet with harvested primary DRGs later incubated in margaric acid-enriched media (Figure 3A). Following confirmation of continued MA enrichment in DRG tissues at the 8-week diet timepoint (Figure 3B), we specifically tested DRG neurons as an established primary cell type commonly used to assess mechanically-evoked currents^20,21,40,41^. Compared with controls, neurons from MA-fed mice displayed an overall reduction in mechanically activated currents across all inactivation types, with statistically significant effects limited to the rapidly inactivating currents. An even more pronounced reduction was observed in cultured DRG neurons exposed to MA-enriched medium (Figure 3C–D). In addition, MA enrichment significantly increased the mechanical displacement threshold required to elicit mechanically gated currents relative to control neurons (Figure 3E). Consistent with these findings, current-clamp recordings during mechanical stimulation revealed a rightward shift in stimulus–response relationships in both margaric acid-enriched groups, indicating elevated mechanical activation thresholds (Figure 3F–G). Together, these results demonstrate that dietary MA supplementation effectively decreases mechanically gated currents in sensory neurons, supporting this lipid manipulation as a feasible *in vivo* approach to inhibit PIEZO channel activity.

**Figure 3.**
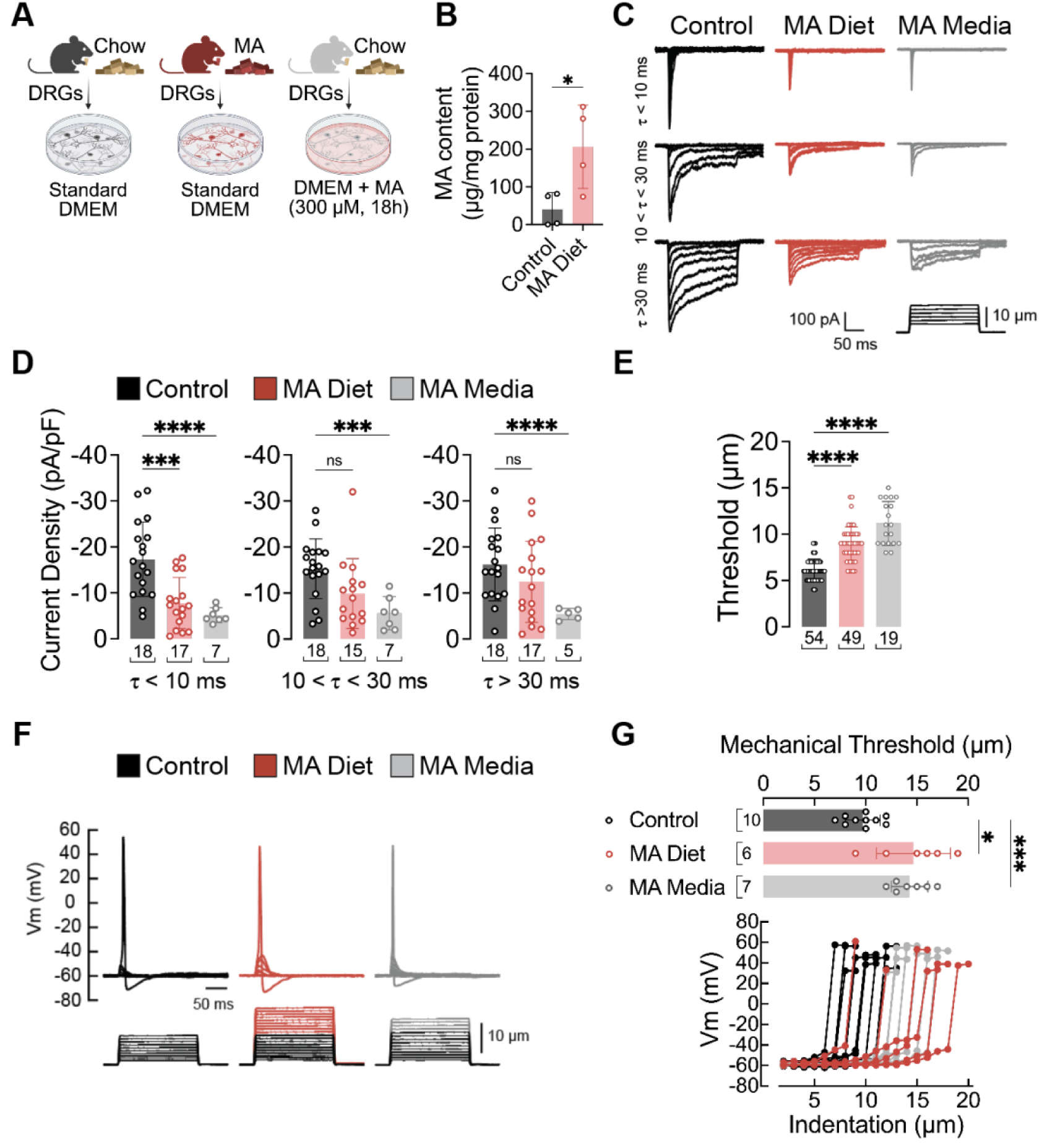
Dietary margaric acid supplementation inhibits mechanically gated currents in DRGs. (A) Illustration showing the experimental design. (B) Liquid chromatography-mass spectrometry analysis of MA content in DRGs tissues from mice fed standard chow diet (Control), Margaric acid-enriched diet (MA Diet), normalized to total protein content. n=4 animals/group. Bars show mean ±SD (error bars), dots show individual animals. Statistical tests: Unpaired *t*-test. (C) Representative whole-cell patch-clamp recording traces showing rapidly-, intermediate-, and slowly-inactivating currents elicited by mechanical stimulation (−60 mV) of DRG sensory neurons from mice fed control diet (*black*), mice fed margaric acid diet (MA diet, *red*), and chow-fed mice DRGs cultured in MA-enriched media (control, *gray*) (300µM, 18h). (D) Current densities elicited by maximum displacement of DRG neurons classified by their time constant of inactivation (τ). (E) Displacement thresholds required to elicit mechano-currents from DRG neurons. (F) Representative current-clamp recordings of membrane potential changes elicited by mechanical stimulation in DRG neurons. (G) Membrane potential peak vs. mechanical indentation of independent mouse DRG neurons. At the top, bar plots show the displacement threshold required to elicit an action potential in these neurons. For C-G bar graphs: bars show mean ±SD (error bars), dots show individual neurons (number of neurons indicated under each bar); animal numbers/group: 5x Control, 5x MA Diet, 4x MA Media; statistical tests: Brown-Forsythe and Welch ANOVA tests with Dunnett’s T3 multiple comparisons test. For all graphs: (ns) *p* > 0.05, (*) *p* ≤ 0.05, (**) *p* ≤ 0.01, (***) *p* ≤ 0.001, (****) *p* ≤ 0.0001.

### Margaric acid-enriched diet reverses urinary dysfunction in aged mice

Given our finding that SMC-specific *Piezo1* deletion protects against aging-related urinary dysfunction, we hypothesized that dietary MA supplementation could improve urinary function in aged mice through its inhibitory effects PIEZO1 channel activity. We assessed urinary behavior in aged mice (∼90 weeks) at baseline while maintained on standard chow and again after margaric acid-enriched diet feeding (4-5 weeks). At baseline, aged mice exhibited the expected urinary dysfunction (Figure 4A–C). Following MA supplementation, these animals showed a significant improvement in urinary behavior, characterized by a reduced voiding frequency and decreased urine deposition in the arena center (Figure 4B, D). This improvement was driven primarily by a reduction in small, secondary voids (Figure 4E–F). Importantly, total urine output was not significantly altered by MA supplementation (Figure 4C), indicating that improved urinary control was not due to a generalized suppression of urination. Although statistically significant changes were detected in other PIEZO-dependent physiological functions, including gait, brush-evoked dynamic touch, and gut function, animals exhibited no changes in von Frey touch sensitivity and no overt health issues (Figure S4). Together, these findings demonstrate that a non-invasive dietary MA supplementation can partially restore urinary control in aged mice.

**Figure 4.**
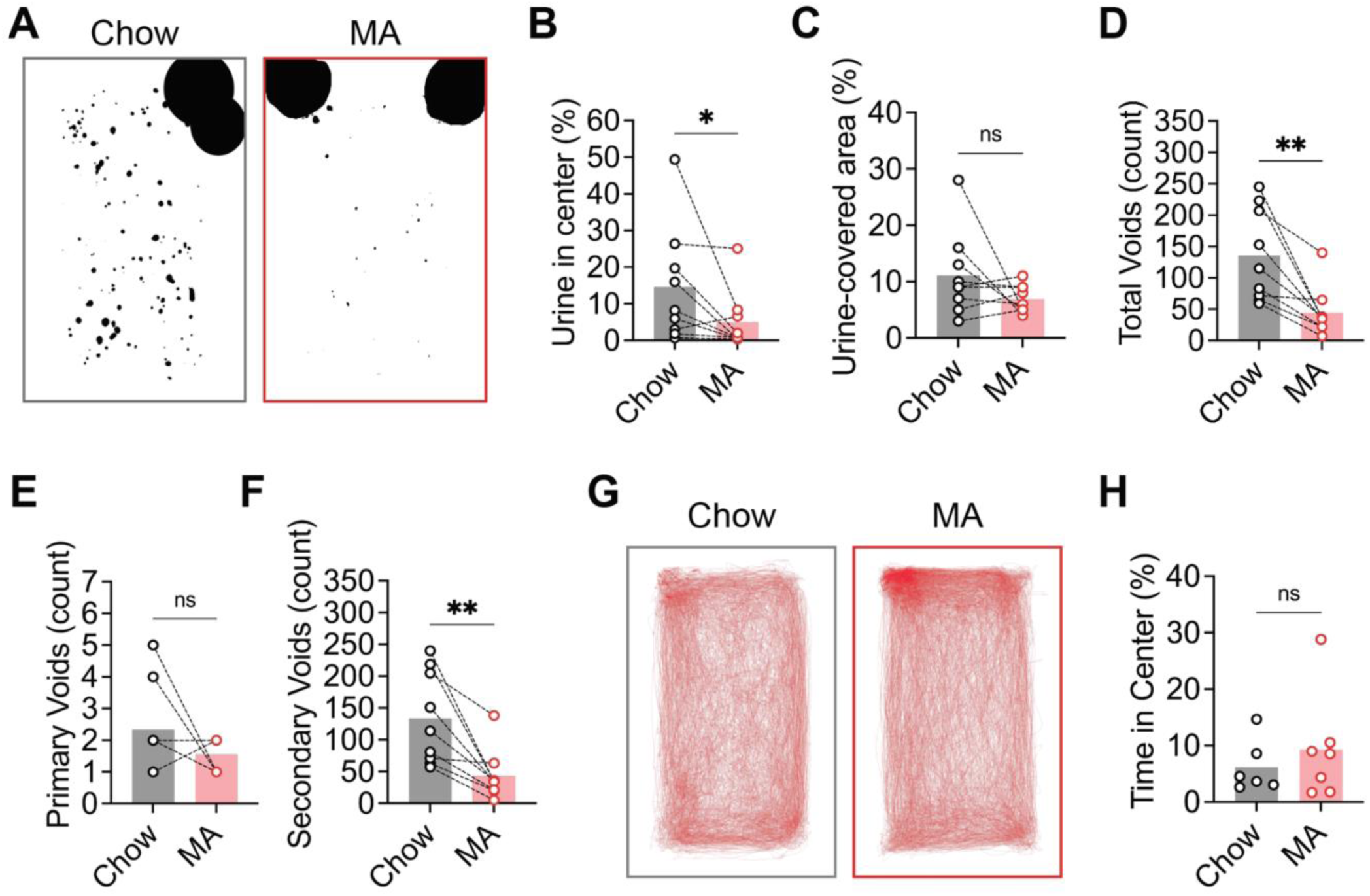
Margaric acid-enriched diet reverses urinary dysfunction in aged mice. (A) Representative images of 4h VSA papers from the same aged mouse tested once while on chow diet (*left, gray*), and again after a 4-5-week margaric acid (MA) diet (*right, red*); urine void spots are shown in black; quantified in D-H (paired analysis). n=9/group. (B) Percentage of urine deposited in center 25% of cage area. n=9/group. (C) Total urine-covered area at the end of a 4 h void spot assay, quantified as percentage of total cage area. n=9/group. (D) Total number of urinary void spots, further stratified by size into primary (E) and secondary (F). n=9/group. (G) Line plots showing representative movement traces from aged chow diet (*left*, gray) and MA diet (*right*, red) mice during the 4h VSA. (H) Time spent in center 25% of cage area quantified; n=6-7/group. For all bar plots: bars show mean, dots show individual animals, and dashed lines indicate paired analysis (when present). Statistical test: (C) Mann-Whitney U test, (D-G) Wilcoxon test, and (H) Paired *t*-test. (ns) *p* > 0.05, (*) *p* ≤ 0.05, (**) *p* ≤ 0.01, (***) *p* ≤ 0.001, (****) *p* ≤ 0.0001.

### Margaric acid rescues aging-related altered bladder pressure-volume dynamics

Aging is known to impair smooth muscle function^29,44,45^, prompting us to investigate its effects on smooth muscle mechanics in the bladder. We isolated the bladder to specifically characterize the contribution of bladder muscle dynamics in the absence of neuronal reflexes. We used an *ex vivo* bladder bath preparation to characterize pressure-volume (P-V) relationships during filling, allowing precise assessment of bladder wall mechanics independent of neuronal reflexes or other systemic influences^46^. The response of the bladder to filling is nonlinear^47^. Normally, detrusor smooth muscle relaxation during bladder filling allows volume expansion while maintaining low pressure until a threshold triggers voiding^48,49^. To examine which phases of the filling/voiding cycle are affected by age and account for differences in bladder size, we parsed the P-V curves into three phases: Phase I, II, and III, corresponding to the initial *passive filling* phase, the *active accommodation* phase (characterized by a pressure plateau or slight drop as the bladder wall relaxes to accommodate incoming fluid), and the post-accommodation phase, marked by a steady pressure rise after active accommodation ends which, *in vivo*, would likely have corresponded to a voiding event^50^.

Aged bladders exhibited markedly altered P-V dynamics compared to young bladders, and after margaric acid-diet, bladders have a P-V profile that is partially restored to that of young animals (Figure 5A-C). Interestingly, the age-related increase in bladder mass with age, a commonly used indicator of bladder remodeling in the setting of dysfunction^51^, was also partially restored toward weights closer to those of young animals (Figure 5D). Specifically, compared to the young chow group, aged chow bladders required larger fill volumes than young chow bladders to reach 15 mmHg, a pressure approximating the typical *in vivo* micturition threshold^52^, consistent with the higher inverse slope (ΔV/ΔP) and increased compliance observed in aged bladders^47^ (Figure 5E-F). Remarkably, aged margaric acid-diet bladders required a significantly smaller fill volume to reach 15 mmHg, which was comparable to the young chow group, and exhibited intermediate average compliance (Figure 5E-F). This overall compliance measurement does not depend on bladder capacity, given that values were normalized to bladder weights. Notably, among the three filling phases, the active accommodation phase (Phase II) in aged chow group was significantly increased relative to the young chow group which showed comparable duration to the aged diet bladders (Figure 5G–I). These results reveal distinct age-dependent differences in volume handling where aged bladders exhibit increased accommodation capacity compared to young bladders. While this increased capacity may allow the bladder to accommodate larger fill volumes as a compensation mechanism for weakened smooth muscle contractility, much like the cardiac smooth muscle, over time, this may lead to a decompensatory phase that mimics myogenic underactive bladder, resulting in decreased voiding efficiency and increased post-void residual volume^53^. Together, these findings reveal a potential mechanism underlying aging-related changes in bladder function.

**Figure 5.**
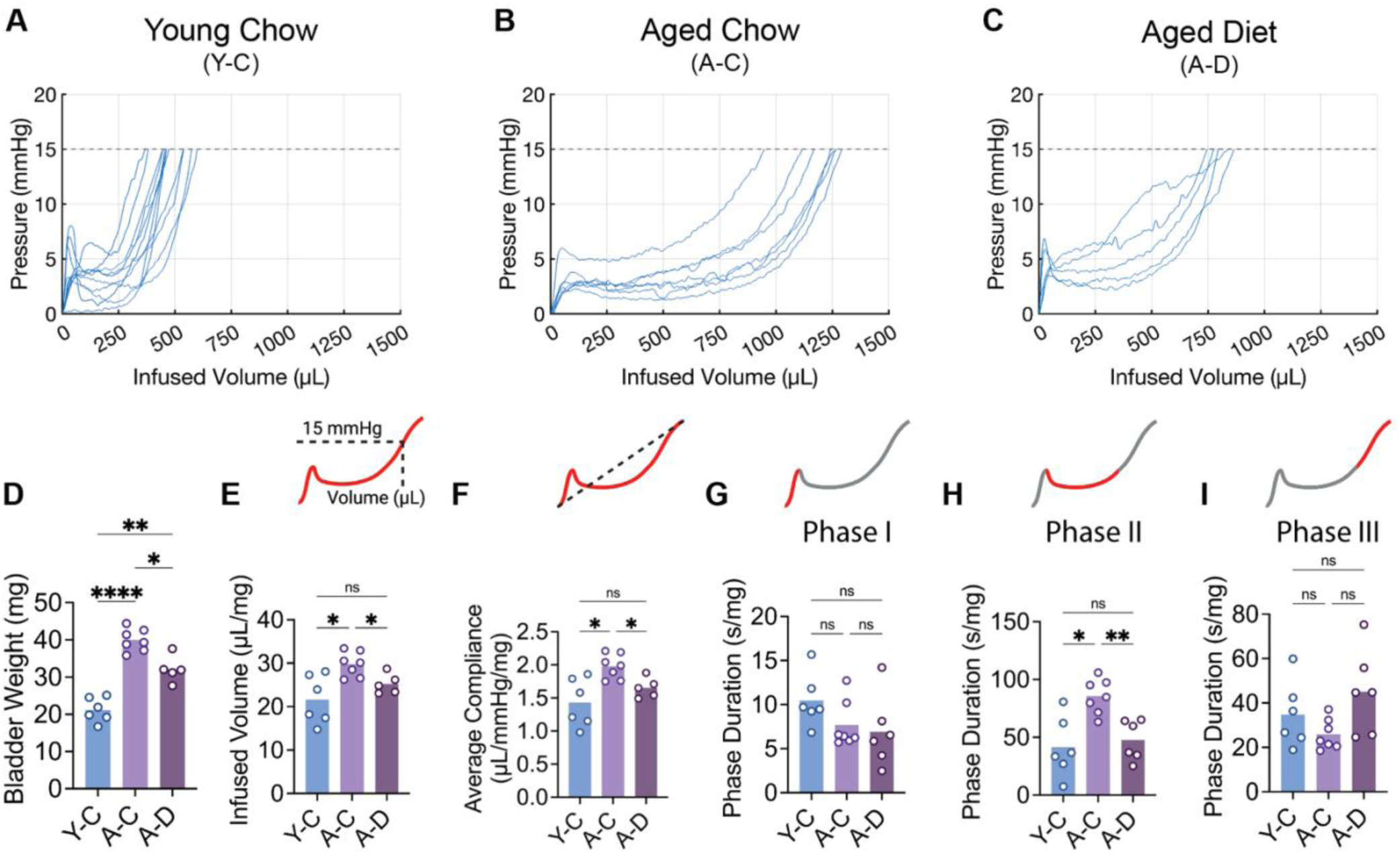
Aging alters bladder pressure-volume handling dynamics which can be partially reversed via dietary inhibition of PIEZO channels. (A-C) Overlayed pressure-volume traces of young chow group (A), aged chow group (B) mice, and aged MA diet group (C) obtained from *ex vivo* bladder pressure-volume assay. Quantified in E-I. n=5-8/group. (D) Bladder weights from young and aged mice. (E) Infused volume needed for bladders to reach voiding pressure point (15 mmHg), normalized to empty bladder weight. (F) Average bladder compliance, calculated as ΔV/ΔP over the 0–15 mmHg filling range of pressure–volume (P–V) curves, normalized to empty bladder weight. (G-I) Duration of each bladder cycle phase, normalized to empty bladder weight. For all bar plots: bars show mean; dots show individual animals. n=5-8/group. Statistical test: Brown-Forsythe and Welch ANOVA tests with Dunnett T3 correction for multiple comparisons. (ns) *p* > 0.05, (*) *p* ≤ 0.05, (**) *p* ≤ 0.01, (***) *p* ≤ 0.001, (****) *p* ≤ 0.0001.

### Ultra-rare and gain-of-function *PIEZO1* variants contribute to bladder dysfunction

Our findings implicating PIEZO1 in urinary dysfunction indicate that variants of this gene could be an underlying risk factor for neuromuscular bladder disorders. Building on evidence that rare germline variants in *PIEZO1* associate with disorders like lymphedema and hereditary xerocytosis^54–58^, we tested whether *PIEZO1* variants contribute to neuromuscular bladder dysfunction. We applied Evolutionary Action (EA)-Pathways analysis^59–63^, an ultra-rare variant, Gene ontology-based (GO)^64,65^ aggregation framework that links genes with specific phenotypes, to United Kingdom Biobank (UKB)^66^ neuromuscular bladder dysfunction samples (cases) and bladder dysfunction-free samples (controls) from aged individuals. EA is a functional impact score derived from the evolutionary importance of the mutated residue (Evolutionary Trace^67^) and the log odds of the observed amino acid change, and it performs well in objective comparisons^68,69^. Among the 29 pathways (i.e., gene sets) recovered from neuromuscular bladder dysfunction cases (Figure 6A), comprising 196 core genes (Table S1), “cation transport” and “detection of mechanical stimulus” were highly significant (Fold-better > 4; reflects the increased EA distribution bias of the pathway relative to same-sized simulated pathways) (Figure 6A). Within the “cation transport” pathway, variants in *PIEZO1/2* were among the core genes driving the EA distribution bias, suggesting a strong correlation to disease (Figure 6B). Interestingly, the *PIEZO1/2*-containing “mechanosensitive ion-channel activity” pathway was also prominent among the 63 pathways recovered from resilient subjects (417 core genes; Table S2). The core bladder dysfunction gene set (cases) and the resilience gene set (controls) overlapped by 20 genes (hypergeometric p = 6.1e-9), including *PIEZO1* and *PIEZO2*, suggesting that divergent phenotypes may arise depending on the specific mutations in these genes.

**Figure 6.**
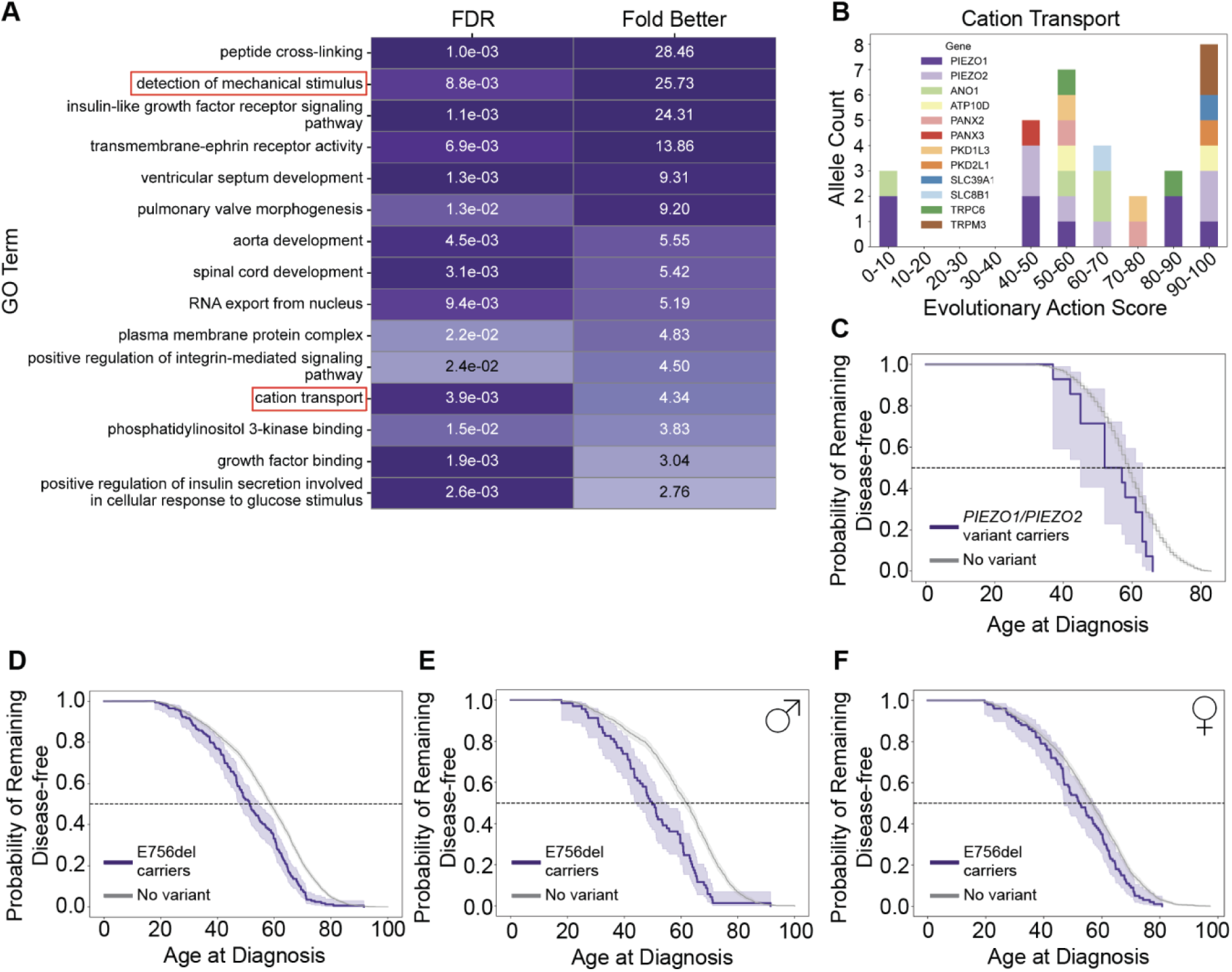
PIEZO1/PIEZO2 variants associated with neuromuscular bladder dysfunction. (A) Pathway-bias associations in UKB cases; “detection of mechanical stimulus” and “cation transport” GO terms highlighted with a red box. (B) PIEZO-centric “cation transport” EA score distribution for UKB cases (B). Lower scores indicate likely benign variants, while higher scores indicate likely high-impact/functionally damaging variants. (C) Kaplan–Meier curve for human subjects from UKB doubleton variants in *PIEZO1/PIEZO2* (n=14 carriers, n=1560 non-carriers; males and females). *PIEZO1*/*PIEZO2* variant carriers, median age: 57, mean age 54.1; No variant, median age: 59, mean age: 58.7. HR = 1.9 (95% CI 1.1–3.2; *p* = 0.02). (D) Kaplan–Meier curve of AoU subjects with disease (N=3191 total) stratified by status as carriers of *PIEZO1* p.E756del (n=169) or non-carriers (n=3022), further stratified by sex: (E) males (n=69 carriers, n=1282 non-carriers) and (F) females (n=100 carriers, n=1740 non-carriers). (D) E756del carriers, median age: 51, mean age 51; No variant, median age: 59, mean age: 57. HR = 1.3 (95% CI 1.1 –1.5; *p* = 5.90e-03). (E) E756del carriers, median age: 50, mean age 50; No variant, median age: 62, mean age: 60. HR = 1.5 (95% CI 1.1 –2.0; *p* = 4.25e-03). (F) E756del carriers, median age: 52, mean age 52; No variant, median age: 57, mean age: 55. HR = 1.2 (95% CI 0.9–1.5; *p* = 0.19).

To assess whether *PIEZO1/2* variants influence clinical features of bladder dysfunction, we stratified cases by individuals with and without an ultra-rare coding variant in *PIEZO1/2* (i.e., doubletons, or variants appearing in no more than two individuals in the UKB) and modelled age at diagnosis. Within the neuromuscular bladder dysfunction group, *PIEZO1/2* variant carriers in the UKB database showed an average of 4.6-year earlier onset (Figure 6C). While these findings show an association, the functional consequences of the variants identified are not known. To determine whether this acceleration in disease onset might be caused by a gain-of-function mechanism, we assessed the well-characterized *PIEZO1* gain-of-function (GOF) indel E756del^15,54^. Given the predominant prevalence of this variant specifically among individuals of African descent^22^, we utilized the NIH *All of Us* (AoU) database^70^, which contains individuals with a broader range of ancestry backgrounds. We found that this variant does not itself strongly associate with neuromuscular bladder dysfunction, where among 3,191 AoU neuromuscular bladder dysfunction cases, 169 (5.3%) were carriers. Although prevalence was unchanged by the presence of this *PIEZO1* GOF variant, likely reflecting the complexity of multifactorial contributions to bladder dysfunction, carriers were diagnosed 7.3-years earlier on average (Figure 6D). Strikingly, this effect was more prominent in males, with carriers being diagnosed 12.2-years earlier on average (Figure 6E), whereas females showed no significant change (Figure 6F). This aligns with our observation that aging-related urinary dysfunction was delayed and more mild in female mice, which corresponds with published work^27,71^ indicating aging-related voiding and storage dysfunction phenotypes are more prominent and appear earlier in male mice (Figures 1 and S1). Collectively, these results indicate that rare, likely GOF *PIEZO1/2* variants accelerate the onset of neuromuscular bladder dysfunction, with the effects being most pronounced in male subjects.

## DISCUSSION

Our findings reveal that the loss of mechanosensory ion channel PIEZO1 in smooth muscle cells (SMCs) protects against aging-related bladder dysfunction, supporting the idea that SMC-specific PIEZO1 contributes to the pathological changes underlying this condition. These results provide new insight into the complex, multifactorial origins of neuromuscular bladder dysfunction during aging. Previous studies examining the neuronal contribution to aging-related urinary dysfunction have produced mixed findings: some report reduced sensory neuron innervation and sensitivity^72,73^ together with impaired reflex function in aged rodents^14,15^ leading to poor bladder-urethral coordination and weaker bladder contractions. These could contribute to urinary retention and potentially leaking or incontinence-like phenotypes. In contrast, other studies have found no differences in innervation, increased innervation, or enhanced sensory neuron sensitivity in aged rodents, underscoring the complexity of the problem^16,17,25,74^. Consistent with this complexity, we observed no alterations in bladder innervation density, although this does not preclude age-related changes in neuronal sensitivity or reflex circuitry. More broadly, alterations in urinary reflexes are inseparable from changes in the bladder itself, where mechanosensory signals originate. While sensitivity to these signals is critical, our reductionist approach was intended to isolate bladder-specific features of aging.

Bladder filling responses are inherently nonlinear because the bladder actively responds to incoming urinary volume. By separating the pressure-volume curve into distinct phases, our analysis captures this nonlinearity and provides a more nuanced view of the bladder’s active accommodation response during filling. Specialized approaches that capture wall thickness and three-dimensional geometry during filling are ideal for fully characterizing functional bladder elasticity^47^. Here, however, we measured thickness and elastic modulus separately. Interestingly, aging increased bladder activity without changing the elastic modulus, which we measured under compression, but overall bladder stiffness will require more sophisticated measurements given age-related changes in wall thickness and size^75^.

Together with our data indicating aged bladders have reduced expression of SMC contractile genes, this suggests that diminished contractile responses lead to increased accommodation during bladder filling with minimal increase in pressure. While such accommodation increases bladder capacity, it likely impairs efficient emptying, predisposing aged animals to incomplete voiding, increased urgency, and incontinence-like phenotypes. Deletion of *Piezo1* from detrusor smooth muscle alters this trajectory. Aged P1KO bladders maintained ATP, EFS, and carbachol responsiveness and showed preserved contractile gene expression, indicating that the reduced PIEZO1-mediated mechanosensation protects intrinsic detrusor contractility. In this context, animals get the beneficial features of increased accommodation capacity combined with relatively preserved contractile function, which may underly the preservation of urinary control in aging.

Multiple structural and cellular properties shape bladder compliance, including wall thickness, cellular composition, extracellular matrix composition, and active smooth muscle response to stretch. Although aging increased wall thickness in control bladders, this remodeling was attenuated in P1KO mice, indicating partial protection from structural changes^76^. The dominant functional effect of *Piezo1* deletion appears to be preservation of smooth muscle contractile capacity, rather than alterations in passive mechanical properties. More work is needed to fully understand the complex molecular, cellular and tissue-level changes that occur with aging and in various disease states^77^. Additionally, how these changes are affecting specific neuronal sensory signals remains unclear, and the mechanosensory reflexes themselves are complex, as they integrate stimuli from bladder stretch and urethral fluid flow. As mechanosensory signals from multiple tissues promote continence and voiding, a more complete picture incorporating multiple tissues will be needed.

It is noteworthy that inhibiting mechanosensory ion channel function through dietary intervention in a mouse that already has age-related urinary dysfunction is sufficient to rescue urinary control and partially restore the pressure-volume handling of aged bladders to a more youthful profile. This could indicate that the dietary inhibition of PIEZOs recapitulates protection from age-related dysfunction seen in *Piezo1* KO animals. Our previous work showed that deleting *Piezo2* leads to an incontinence-like phenotype similar to that seen in aging^6^, so we would expect that significantly inhibiting PIEZO2 (broadly in caudal central nervous system tissues) should exacerbate urinary dysfunction. Although membrane margaric acid enrichment has been shown to modulate both PIEZO1 and PIEZO2, it exerts a stronger inhibitory effect on PIEZO1 activity^21^. We therefore interpret the beneficial effects of the MA diet on age-related urinary dysfunction as arising primarily through PIEZO1 in bladder smooth muscle, while not excluding contributions from other mechanosensory tissues within the bladder. Our results are consistent with successful human therapies for conditions like overactive bladder that also target the detrusor muscle neurotransmitter pathways, but their broad reach to other smooth muscle organs bring many unwelcome side-effects^78^. This work offers a potential for novel therapeutic avenue by instead targeting mechanosensory pathways in the detrusor muscle.

Aging-related bladder dysfunction exhibits notable sex differences^7,27,52,71^, although interpretation of human data is complicated by sex-specific anatomical and physiological factors. In men, lower urinary tract dysfunction in later life is frequently influenced by prostate enlargement and bladder outlet obstruction, which can secondarily drive or exacerbate bladder neuromuscular dysfunction^79–81^. In women, lifetime exposures such as pregnancy and parturition, together with hormonal factors, contribute to bladder structural and functional changes with aging^82–84^. Fluctuations in sex hormones are known to influence bladder smooth muscle, urothelial signaling, and neural control, and the hormonal landscape diverges markedly between sexes with age, most prominently with menopause in women^85^. Previous work found muscle strength was not altered, but compliance was higher in aged female mice^5^. Notably, our study assessed only *nulligravida* (never pregnant) female mice, limiting the ability to model the combined effects of reproductive history and age-related hormonal transitions observed in the human female population, which may partially account for differences in incidence and prevalence between clinical and experimental settings. Despite this, male mice exhibited a more robust bladder dysfunction phenotype with aging compared to females, suggesting an intrinsic sex difference not attributable to reproductive history^86^. Interestingly, this pattern parallels observations in humans, where males with neuromuscular bladder dysfunction are diagnosed present at a significantly younger age when carrying a *PIEZO1* gain-of-function variant, implicating enhanced mechanotransduction, potentially modulated by sex-specific factors, as a contributor to earlier disease onset and severity. Together, these findings support a model in which sex-specific hormonal, mechanical, and neuromuscular factors interact to shape susceptibility to aging-related bladder dysfunction, with PIEZO1-mediated mechanotransduction contributing via the muscular axis.

## SUPPLEMENTARY FIGURES

**Figure S1.**
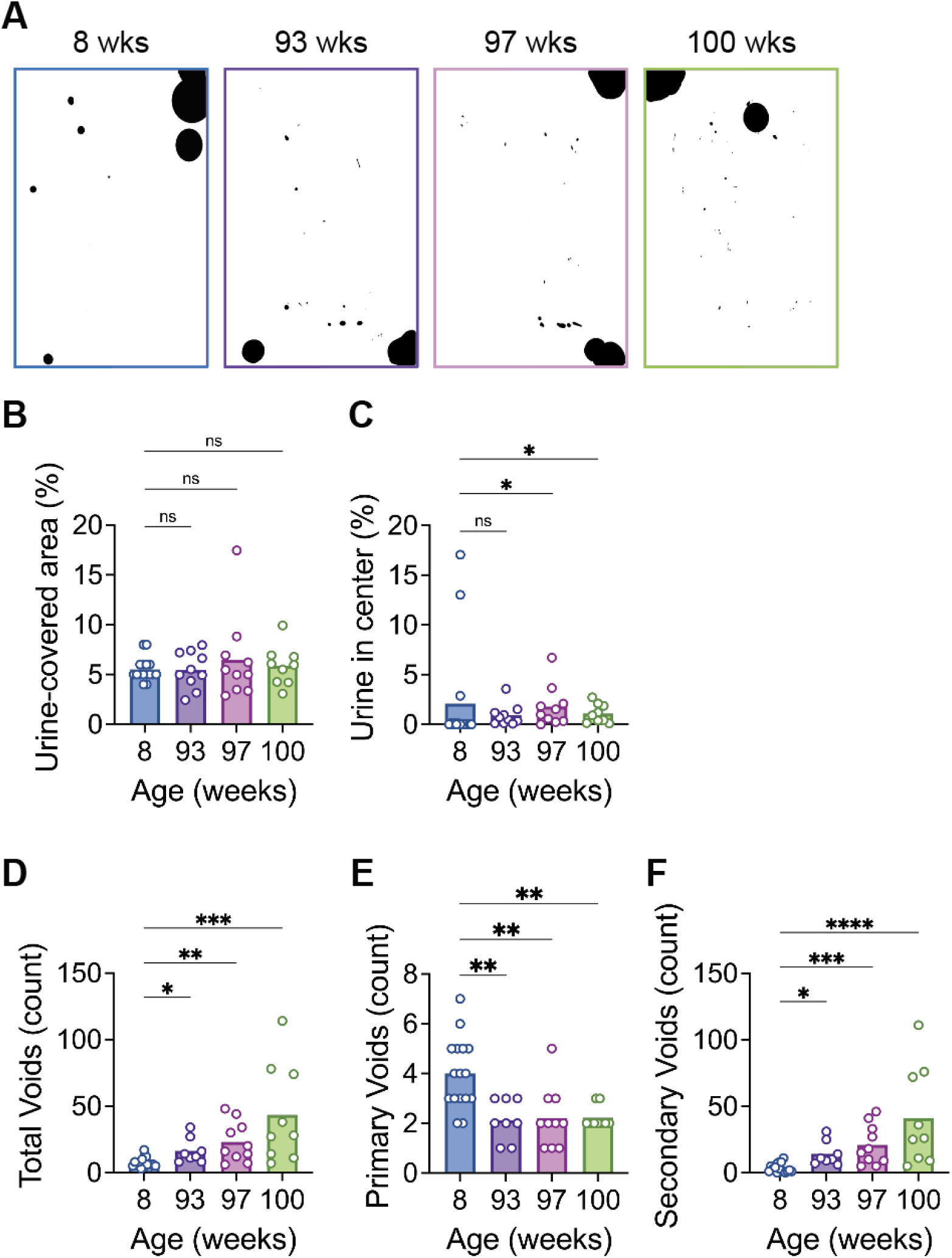
Aging-related changes in urinary behavior are more subtle in aged female mice than male mice. (A) Representative images of 4h VSA papers from young (8-week-old) and aged (93-, 97-, and 100-week-old) female mice. Urine void spots are shown in black. (B) Total urine-covered area at the end of a 4 h void spot assay, quantified as percentage of total cage area; n=9-16/group. (C) Percentage of urine deposited in center 25% of cage area; n=9-16/group. (D) Total number of urinary void spots, further stratified by size into primary (E) and secondary (F); n=9-16/group. For all bar plots: bars show mean; dots show individual animals. Statistical test: (B-F) Kruskal-Wallis test with Dunn’s multiple comparisons ad hoc test. (ns) *p* > 0.05, (*) *p* ≤ 0.05, (**) *p* ≤ 0.01, (***) *p* ≤ 0.001, (****) *p* ≤ 0.0001.

**Figure S2.**
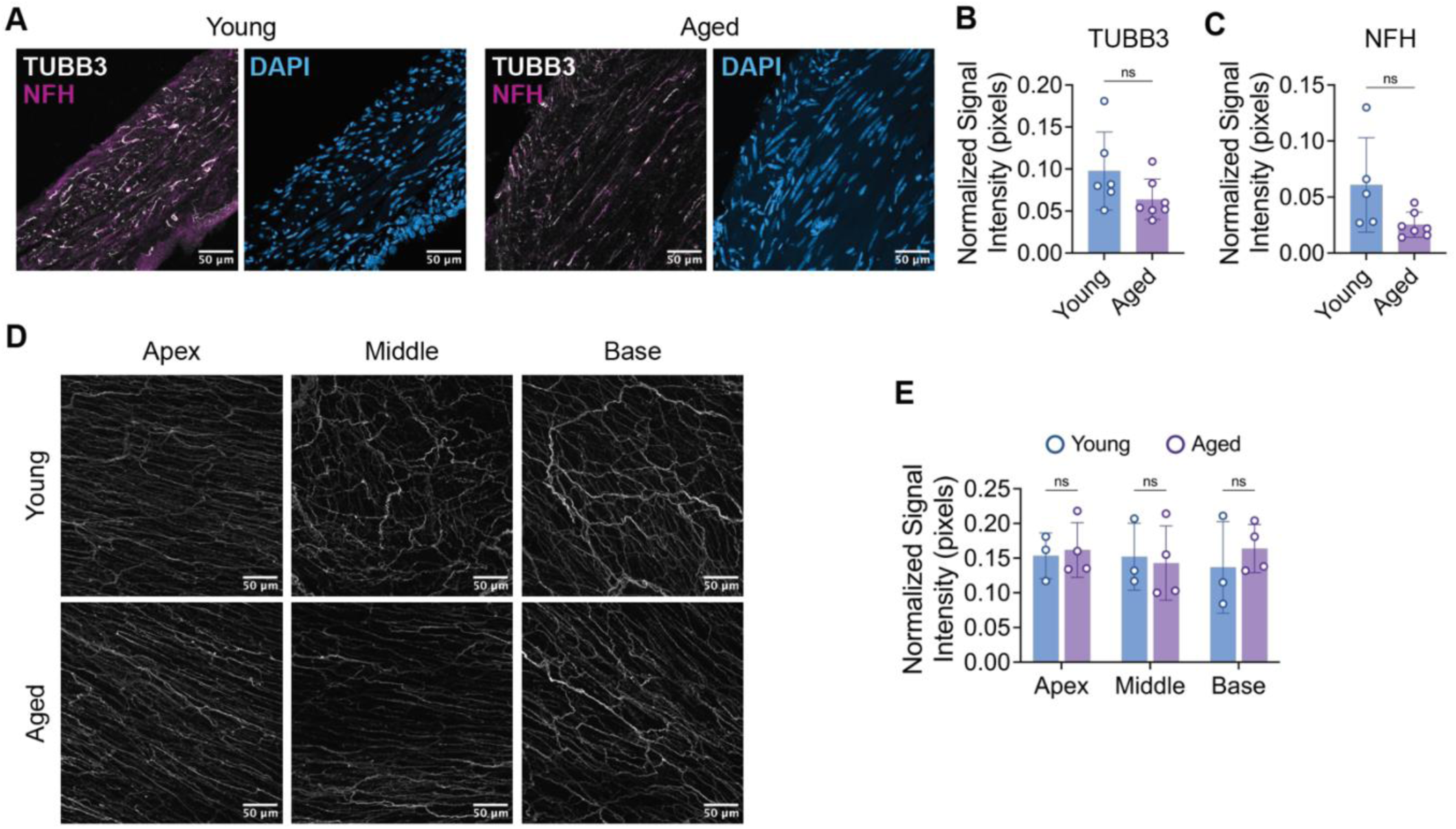
Bladder muscle innervation is similar in young and aged mice. (A) Representative immunohistochemistry images from young and aged mouse bladders tissues sections, labelled against TUBB3 (pan-neuronal marker) and NFH (myelinated neurons). (B-C) Quantification of tissue section immunohistochemistry, shown as signal pixel intensity normalized to tissue area; n=6-7/group. (D) Representative immunostaining images from whole-mount young and aged mouse bladders labelled against TUBB3 (pan-neuronal marker). (E) Quantification of whole-mount immunohistochemistry, shown as signal pixel intensity normalized to tissue area; n=3-4/group. For all bar graphs: bars show mean ±SD (error bars), dots show individual animal means. Statistical tests: (B-C) Welch’s *t*-test. (E) Two-way ANOVA with Šídák’s multiple comparisons test. (ns) *p* > 0.05, (*) *p* ≤ 0.05, (**) *p* ≤ 0.01, (***) *p* ≤ 0.001, (****) *p* ≤ 0.0001.

**Figure S3.**
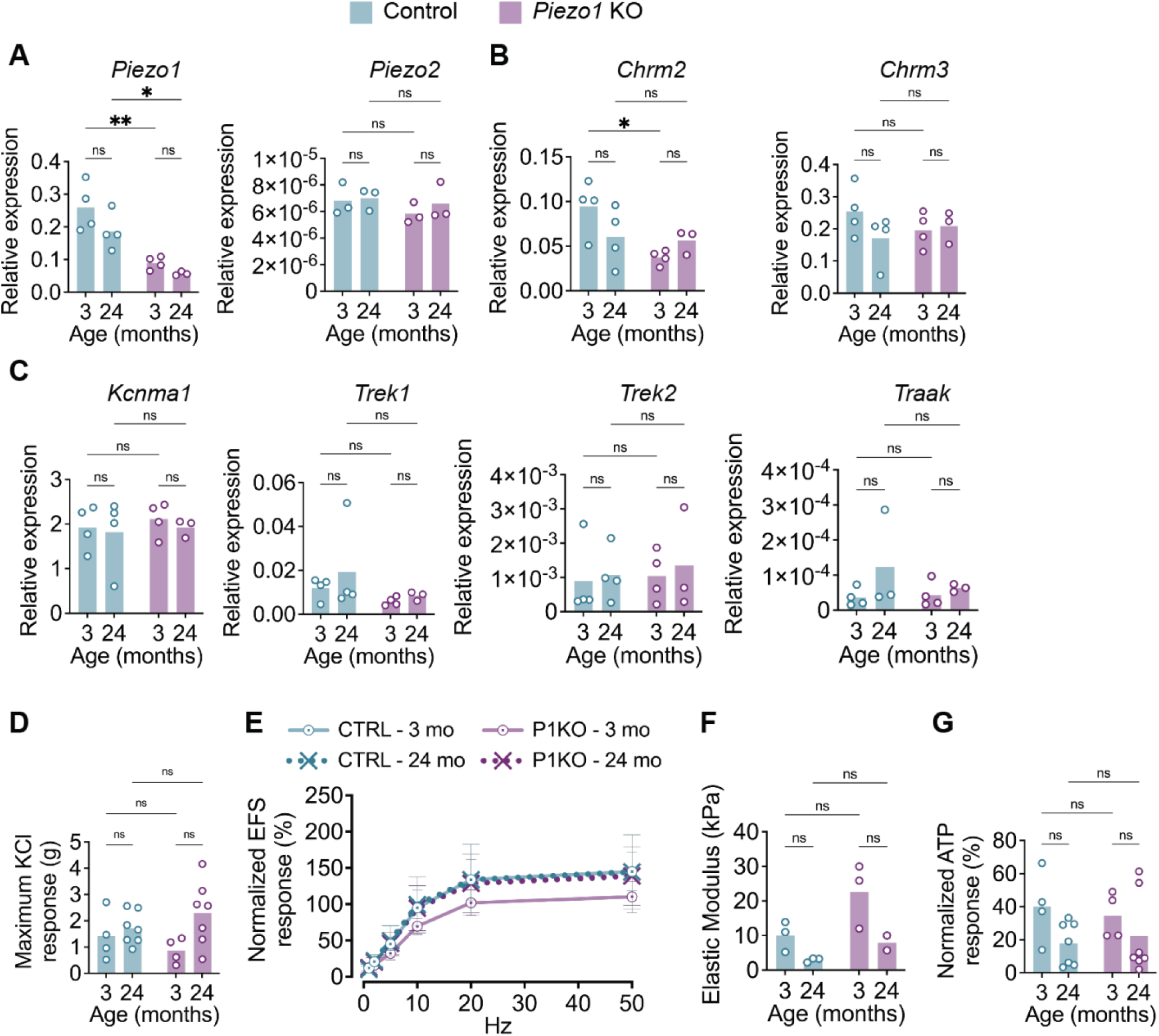
Smooth muscle-specific deletion of *Piezo1* alters bladder muscle properties. (A-C) Relative mRNA expression levels of select genes in bladder smooth muscle tissues, normalized to reference genes. (A) *Piezo1* and *Piezo2* ion channels; (B) muscarinic receptors; (C) major potassium channels. n=3-4/group. (D) KCl-induced maximal bladder muscle contractile response; n=4-7/group. (E) Bladder muscle contractile response to electric field stimulus (EFS) at frequencies of 1, 2, 5, 10, 20, and 50 Hz; normalized to KCl-induced maximal response; n=4-7/group. (F) Elastic modulus of isolated bladder muscle strips; n=2-3/group. (G) Bladder muscle contractile response to 1 mM ATP, normalized to KCl-induced maximal response; n=4-7/group. For all bar graphs: bars show mean; dots show individual animals; for (E): dots show means ±SD (error bars). Statistical tests: (A-D, F-G) Two-way ANOVA with Šídák’s multiple comparisons test; (E) Two-way mixed-effects model (REML) with Tukey’s multiple comparisons test; no significance bars = not significant. (ns) *p* > 0.05, (*) *p* ≤ 0.05, (**) *p* ≤ 0.01, (***) *p* ≤ 0.001, (****) *p* ≤ 0.0001.

**Figure S4.**
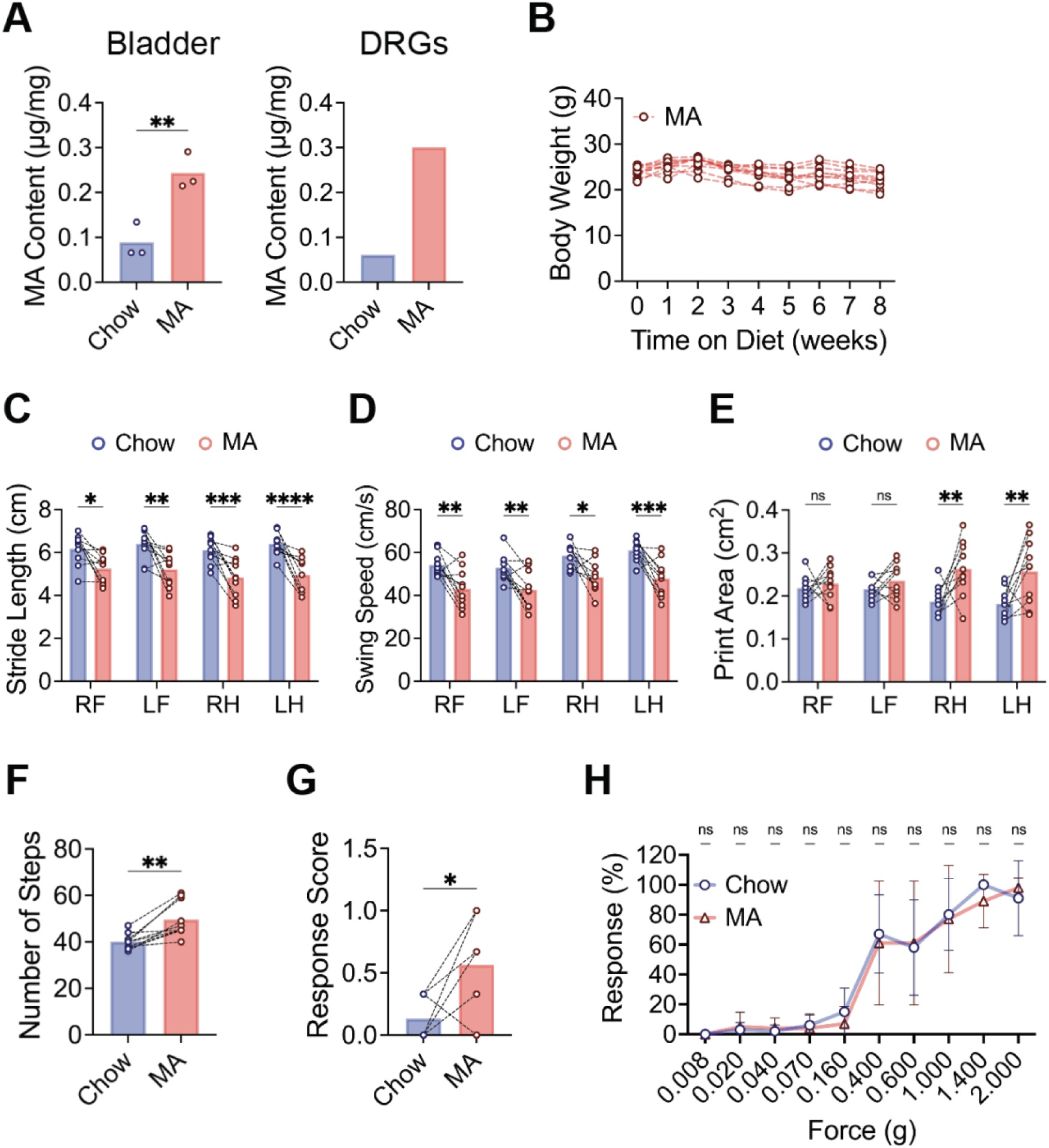
Characterization of systemic effects of dietary PIEZO channel manipulation in young, healthy mice. (A) Liquid chromatography-mass spectrometry analysis of margaric acid (MA) content in bladder (*left*) and DRGs tissues (*right*) from young mice fed standard chow diet (Chow, *grey*) or MA-enriched diet (MA, *red*). n=3/group; for DRG tissues, samples from 3 animals/group were pooled; bars show mean, dots represent individual animals. Statistical test for bladder tissues: Welch’s *t*-test; statistical analysis for DRGs could not be performed since all 3 samples/group were pooled to meet minimum tissue weight for analysis. (B) Chart showing body weight over 8 weeks for young mice fed MA diet. Week 0 indicates weight while on standard chow prior to the start of MA diet feeding. Individual animals shown. (C-F) CatWalk gait analysis of young mice fed standard chow or MA diet. (C-E) Two-way repeated-measure ANOVA with Šídák’s multiple comparisons test; (F-G) Paired *t*-test. (G) Dynamic touch (brush) response scores for young mice tested at baseline (chow diet) and again after MA diet. Paired *t*-test. (H) Von Frey touch sensitivity assay showing percentage response to each force in young mice fed chow or MA diet. Dots show mean ±SD (error bars). Two-way repeated-measure ANOVA with Šídák’s multiple comparisons test. For all bar plots: bars represent mean; dots represent individual animals. (B-I) n = 10/group. (ns) *p* > 0.05, (*) *p* ≤ 0.05, (**) *p* ≤ 0.01, (***) *p* ≤ 0.001, (****) *p* ≤ 0.0001.

## METHODS

### EXPERIMENTAL MODEL AND STUDY PARTICIPANT DETAILS

#### Mice

Aged (90 weeks), wild-type C57BL/6J male and female animals were purchased from The Jackson Laboratories and housed in Baylor College of Medicine’s clean Transgenic Mouse Facility (TMF) in accordance with the approved IACUC guidelines (14 h/10 h light/dark cycle at 21 °C with 40-70% humidity). Mice were allowed at least one week to acclimate to their new housing in TMF before any testing was performed. All aged mice of the same sex were housed in pairs and continued to receive ad libitum water and irradiated PicoLab® Select Rodent 50 IF/6F rodent diet (LabDiet, #5V5RZ), until the start of the diet experiments.

To obtain SMC specific *Piezo1* knockout (P1KO) mouse model [*Myh11-creERT2; Piezo1^fl/fl^*], we *crossed Piezo1 flox B6.Cg-Piezo1^tm^*^2^*^.1Apat/J^*mouse (The Jackson Laboratories, strain #029213; gift from Dr. Fouad Chebib, Mayo Clinic)^87^ *with B6.FVB-T (Myh11-cre/ERT2)1Soff/J* (The Jackson Laboratories, strain #019079)^88^. Since the Cre-carrying BAC transgene in the *Myh11-creERT2* line was inserted on the Y chromosome^88^, only male carriers could be produced. P1KO male mice received a gavage of tamoxifen (Sigma, Cat. #T5648-1G) (150 mg/kg in 0.2 mL corn oil) for three days at 7-8 weeks to induce SMC-specific *Piezo1* KO. Mice were euthanized by CO_2_ inhalation followed by cervical dislocation at appropriate age timepoints. All animals were housed in the Mayo Clinic animal facility (12 h/12 h light/dark cycle) and experimental procedures were approved by the Institutional Animal Care and Use Committee.

For electrophysiology experiments (Figure 3), mice procedures described were reviewed and approved by the University of Tennessee Health Science Center (UTHSC) Institutional Animal Care and Use Committee (IACUC). All methods were carried out in accordance with approved guidelines. Adult (2- to 4-month-old) wildtype male C57BL/6J mice were obtained from The Jackson Laboratory (Stock No. 000664) and housed in a 12 h/12 h light/dark cycle at 21 °C with 40-60% humidity. Food (standard chow) and water were provided ad libitum, unless otherwise stated in specific methods.

## Human subjects

### Selection of bladder dysfunction cohorts from the UKB and AoU

#### UKB Cohort

We extracted bladder dysfunction cases from the UKB using the following criteria: White ethnic background (f.21000); “Neuromuscular dysfunction of bladder, not elsewhere classified” ICD10 diagnoses (N31 series; f.41270); no history of multiple sclerosis (G35 series; f.41270) and chemotherapy (Z512, Z926, Z511). We selected bladder dysfunction-resistant controls using the following filters: White ethnic background (f.21000); no history of bladder dysfunction (N31, N32 series; f.41270), neurological disorders (G10-G37 series), diabetes (E11-E14), and bladder cancer (C67 series); no history of neurological disorders or diabetes in mother, father, and siblings (f.20107, f.20110, f.20111); a normal body mass index (BMI) at intake (18.5-25; f.21001); current age > 75 (70% of cases were diagnosed by 75). Bladder dysfunction cases and resistant controls were subjected to sample-level quality control (QC) and stratified by sex, resulting in 1,574 cases and 1,574 controls. Phenotype field instances used for cohort selection were downloaded in April 2024.

#### AoU Cohort

We extracted genotypes and phenotypes from samples with short-read whole genome sequencing and electronic health record (EHR) data (n = 340,582) within the Controlled Tier v8 dataset (accessed May 2025). Bladder dysfunction cases were selected using the following criteria: “White” race; “not Hispanic or Latino” ethnicity; presence of N31 series ICD10 code in the “Condition” domain; no multiple sclerosis (G35 series) ICD10 code in the “Condition” domain; no “Chemotherapy” keyword in the “Procedures” domain. Bladder dysfunction-resistant controls were selected using the following criteria: “White” race; “not Hispanic or Latino” ethnicity; normal BMI (18.5-25) in the “Physical Measurements” domain; no presence of bladder dysfunction/urinary system conditions (N31, N32, R32 N39, and R39 series), neurological disorders (G10-G37 series), diabetes (E11-E14 series) and bladder cancer (C67 series) ICD10 codes in the “Condition” domain; current age > 75; no family history, defined in the Survey>Personal and Family Health History series, of prediabetes, type 1 diabetes, type 2 diabetes, other diabetes, cerebral palsy, dementia (Alzheimer, vascular, etc.), epilepsy/seizure, Lou Gehrig’s disease, memory loss or impairment, multiple sclerosis, muscular dystrophy, Parkinson’s disease, and other brain or nervous system conditions, using the survey question “Including yourself, who in your family has had [condition]? Select all that apply.” We subjected bladder dysfunction cases and resistant controls to sample-level QC and subsequently selected a cohort, stratified by sex, of 1,431 cases and 1,431 controls.

#### AoU PIEZO1:E756del Cohort

We identified 3,191 samples diagnosed with “neuromuscular dysfunction of the bladder, not elsewhere classified” (N31 ICD10 series). Of these, 169 harbored at least one copy of the PIEZO1:p.E756del allele, a known gain-of-function indel^22,89^ (AoU v8 dataset, accessed May 2025).

## Treatments

### Diet supplementation

For electrophysiology experiments (Figure 3), wildtype male C57BL/6J mice (6-8 weeks old) were pair-fed for 8 weeks (before DRG dissection) with a: A) control diet, modified AIN-93G purified rodent diet (Dyets, #105012) with 59% fat-derived calories from anhydrous milk fat (kcal/kg): casein (716), L-cystine (12), maltose dextrin (502), cornstarch (818.76), anhydrous milk fat (2430), soybean oil (630), mineral mix #210025 (30.8), and vitamin mix #310025 (38.7), or B) an isocaloric margaric acid enriched diet, modified AIN-93G purified rodent diet (Dyets, #104839) with the same components except for the 59% fat-derived calories being from margaric acid (heptadecanoic acid).

For physiological/behavior assays (Figure S4), wildtype male C57BL/6J mice (9-10 weeks old) were pair-fed for 6 weeks with a: A) control linoleic acid enriched diet, (Dyets, #105182) with 59% fat-derived calories from safflower oil (kcal/kg): casein (716), L-cystine (12), maltose dextrin (882), cornstarch (458.8), safflower oil (2430), soybean oil (630), mineral mix #210025 (30.8), and vitamin mix #310025 (38.7), or B) an isocaloric margaric acid enriched diet, modified AIN-93G purified rodent diet (Dyets, #104839), with 59% fat-derived calories from margaric acid (heptadecanoic acid). Aside from the main source of fat, both diets have identical ingredients with slightly modified composition to enable comparable formulation; see description of margaric acid enriched diet in the above paragraph. The only exception was Figure S4A where young, wild-type females were used instead (9-19 weeks).

## Histology

### Immunohistochemistry

Young and aged male and female mice were euthanized according to the lab’s IACUC-approved protocols, perfused with ice-cold 20-25 mL 1X phosphate buffered saline (PBS) followed by 20-25 mL ice-cold 4% paraformaldehyde (PFA, diluted in 1X PBS). Bladders were dissected, emptied of urine, then briefly washed in 1X ice-cold PBS before being drop-fixed in 4% PFA overnight at 4°C (Thermo Scientific, 047377.9M). Tissues were then either directed to whole-mount staining or the tissue section staining (see protocols below).

**For tissue sections:** tissues were moved from 4% PFA to 30% sucrose in 1X PBS (Sigma-Aldrich, S0389-500G) for cryopreservation and incubated overnight at 4°C. Bladders were then embedded in Tissue-Tek® O.C.T. Compound (Sakura Finetek, 4583), and flash frozen in liquid nitrogen. Fixed tissues were then sectioned at 30 µm and collected onto Superfrost™ Plus Microscope Slides (Fisher Scientific, 12-550-15). Slides were washed with 1X PBS for 5 min. on rocker. A hydrophobic barrier was drawn around tissues using ImmEdge® Pen (Vector Laboratories, H-4000), then tissues were blocked with 5% blocking solution, made of 5% goat serum (Abcam, ab7481) in 0.3% PBST (0.3% triton X in 1X PBS), for 1h at room temperature (RT). Slides were co-stained using rabbit anti-B3-Tubulin (diluted 1:1000 in blocking solution, Abcam, ab18207) and chicken anti-NFH (Abcam, ab4680) overnight at 4°C. At RT, slides were washed with 1X PBS for 3 x 5 min. on rocker then incubated for 1h with secondary antibodies diluted 1:1000 in blocking solution (Goat anti-rabbit 594, ab150080, Abcam and goat anti-chicken 488, Abcam, ab150169). Slides were washed in 1X PBS for 5 x 10 min. at RT. Finally, tissue slides were mounted using ProLong™ Gold Antifade Mountant (ThermoFisher Scientific, P10144) and imaged under a 20X objective on a Zeiss LSM 900 confocal microscope.

**For whole-mounts:** at room temperature (RT), fixed bladders were then washed for 5 min. in 1X PBS, 15 min. in fresh 1X PBS, then washed for 1h in 0.3% PBST (0.3% triton X in 1X PBS) on rocker. Bladders were then moved to fresh 0.3% PBST (0.3% triton X in 1X PBS) and washed overnight on rocker (RT). Bladders were incubated in rabbit anti-beta III Tubulin primary anti-body solution (ab18207, Abcam; diluted 1:500 in blocking solution) for 5 days at 4°C on rocker. Following primary anti-body incubation, bladders were washed in 0.3% PBST (0.3% triton X in 1X PBS) 2 x 1h on rocker, then washed in fresh 0.3% PBST (0.3% triton X in 1X PBS) overnight at RT. Finally, tissues were incubated in goat anti-rabbit IgG H&L Alexa Fluor® 594 antibody solution (ab150080, Abcam; diluted 1:500 in blocking solution) for 24h at 4°C on rocker (dark). Stained bladders were washed 5 x 1h in fresh 1X PBS on rocker at RT (dark), then cleared overnight in EasyIndex (EI-500-1.52, life canvas technologies). For imaging, bladders were mounted with EasyIndex between two coverslips (taped), allowing imaging from both sides. Mounted bladders were imaged at 10X and 20X using a confocal microscope (LSM900, Zeiss), capturing the base, middle, and apex areas.

### Quantification of nerve density in bladder

A standardized number of images (5 for tissue sections, 3 for whole-mounts) were quantified per bladder per marker. Raw CZI files were imported into ImageJ/FIJI and viewed as hyperstacks in composite color mode. Z-stacks were generated using maximum-intensity projection, with the number of optical slices standardized across all images to ensure consistency. Images were converted to 8-bit format prior to thresholding, and individual channels were separated for analysis. Regions of interest (ROIs) corresponding to the tissue area were manually delineated using the polygon selection tool and stored in the ROI Manager. The channel of interest was thresholded using a standardized threshold range across all images (binary conversion from standardized threshold to 0, indicating background, and 255, indicating signal) to minimize background and false-positive signal. Signal quantification (in pixels) was performed by analyzing the histogram of the thresholded binary image within the ROI. Innervation density was calculated as the number of threshold-positive pixels (intensity = 255) divided by the total ROI area (sum of signal and background pixels), accounting for differences in tissue size across images.

## Tissue functional assays

### Bladder stiffness and thickness measurement

Freshly harvested full thickness bladder was cut open and placed on a glass slide as flat sheet. The stiffness and thickness of the tissues were tested using a spherical indenter (radius = 0.25 mm) in a bath of heated PBS (37 °C ± 1 °C) with the MicroTester G2 (CellScale, ON, Canada). Tissues were indented to 10% of the tissue thickness in 30 seconds, held at that depth of 5 seconds, and then unloaded in 30 seconds. The loading component of each trial was fitted with the Hertz contact model^90^.

### Bladder bath analyses

Bladder bath assays were performed as described in previous studies^91,92^. Mice were euthanized via CO_2_ inhalation, and the bladders were immediately excised. Whole bladders were transported overnight on wet ice in Belzer UW Cold Storage Solution (Thermo Fisher Scientific, # NC2122383)^93^ to the laboratory in UW–Madison. Upon arrival, bladders were bisected longitudinally, and each half was secured with 5-0 silk sutures (Fine Science Tools, Cat. #1802050) at both ends. The tissues were then mounted between an arm and a force transducer (Grass Instruments, FT-03). Contractile responses were recorded as changes in isometric tension (g).

The tissues were submerged in a 37 °C, water-jacketed tissue chamber containing Krebs solution (133 mM NaCl, 16 mM NaHCO_3_, 5 mM KCl, 1 mM MgCl_2_, 1.4 mM NaH_2_PO_4_, 2.5 mM CaCl_2_·2H_2_O, 7.8 mM d-glucose, pH 7.2) aerated with a 95% O2 and 5% CO_2_ gas mix. Bladder tissues were maintained at a baseline tension of 1 g for 60 minutes before starting experiments, with the Krebs solution refreshed every 15 minutes. Electrical field stimulation (EFS) was conducted by stimulating the bladders with 10 V, 0.5 ms pulse duration, at frequencies of 1, 2, 5, 10, 20, and 50 Hz (Grass Instruments, Cat. #S48). Each frequency was applied for 5 seconds, with a 3-minute recovery period between stimulations.

Following EFS, bladders were allowed to recover for 30 minutes, with Krebs solution refreshed after 15 minutes and tension adjusted to 1 g as needed. Carbachol (Thermo Fisher Scientific, AAL0667403) was then added to the baths in increasing concentrations (0-10 µM final concentration, from 1000× fresh stocks), with each addition made once the response to the previous dose plateaued, to induce a stepwise contractile response.

After the final carbachol application, bladders were washed three times with Krebs solution, and tension was readjusted to 1 g as needed. A 30-minute recovery period followed, with Krebs changed at the 15-minute mark. ATP (Sigma, A1852-1VL) was then applied (1 mM final concentration, from 1000× fresh stock).

After the final ATP application, the bladders were washed three times with Krebs, and tension was readjusted to 1 g as necessary. Following a 30-minute recovery period, 60 mM KCl was applied to the baths to elicit a maximal contractile response. Baseline-subtracted responses were reported as “Maximum KCl response”.

For all bladder bath analyses, bladder contractility was recorded and analyzed using Labchart software (ADInstruments, Cat. #v8.1.30) by an investigator blinded to treatment conditions. Responses were baseline-subtracted, normalized to KCl-induced maximum response, and expressed as a percentage. If a bladder half failed to respond to KCl or EFS, it was excluded from all bladder bath analyses

### Ex vivo bladder pressure-volume analysis

A mouse ex vivo bladder prep was set-up, as described in (Durnin *et al.*, 2018)^46^. Briefly, for this study we utilized the gastrointestinal motility set up from Med Associates Inc. A small organ bath with tubing kit (Med Associates Inc., GIM-101 and TBU-GIMM-SUPERFUSE) was continuously perfused with circulating Krebs–bicarbonate solution (KBS, Thermo Scientific, J67591.K2) solution (maintained at 37°C, oxygenated with 95% O_2_/5% CO_2_ gas mix) using a gas dispersion tube (Med Associates Inc., GD-L). Continuous circulation was achieved using a digital MasterFlex® L/S® Digital Miniflex® single-channel peristaltic pump (Avantor, 07525-40). Briefly, mice were euthanized in accordance with the lab’s approved IACUC protocols, perfused with ice-cold 1X PBS, then the bladders with intact ureters and urethra were immediately collected in pre-oxygenated KBS solution in a black Pyrex bottom Sylgard dissection dish where they were cleaned from attached connective tissues and emptied of urine by gently pressing on the bladder in the apex to urethra direction on a Kimwipe. Both ureters were then tied off using a 6-0 silk suture (Roboz Surgical Instrument Co., SUT-14-1) and bladders were transferred to the organ bath. Bladders were then catheterized with a blunt 25G, 0,5” needle (SAI Infusion Technologies, B25-50) connected to a single-channel syringe pump (Braintree Scientific Inc., BS-8000), filling the bladder with oxygenated KBS at 15 µL/min. Syringe pump tube was also connected through a three-way stopcock to a pressure sensor (BIOPAC Systems Inc., RX104A), which records pressure through a pressure transducer and general purpose transducer amplifier system (BIOPAC Systems Inc., TSD104A and DA100C). This set-up allows live recording of intravesicular bladder pressure in response to controlled volume filling using a data acquisition system with AcqKnowledge software (BIOPAC Systems Inc., MP160WSW). Finally, pressure-volume curves were analyzed using a custom-made MATLAB code by an experimenter blinded to the animals’ information.

## Molecular Assays

### Reverse Transcriptase Quantitative Polymerase Chain Reaction (RT-qPCR)

RNA was isolated according to RNeasy MicroKit (Qiagen, Cat. #74134) instructions. RNA reverse transcription was completed using SuperScript VILO cDNA Synthesis Kit (Invitrogen, Cat. #11754050) and a PCR of 10 min at 25 °C, a 60 min. cycle at 42 °C, and 5 min at 85 °C. cDNA was diluted and analyzed for *Smtn, Myh11, Acta2, Piezo1, Piezo2, Chrm2, Chrm3, Kcnma1, Trek1, Trek2,* and *Traak* (Table 1). All experiments were done by reverse transcriptase quantitative polymerase chain reaction according to LightCycler 480 SYBR Green Master I (Roche, Cat. #04707516001) instructions on a LightCycler® 480 II, 96 (well) system (Roche, Cat. #05015278001). Expression values were normalized to the geometric mean of reference genes (*Gapdh, Actb,* and *Hprt*) using the ΔCt method.

**Table 1:**
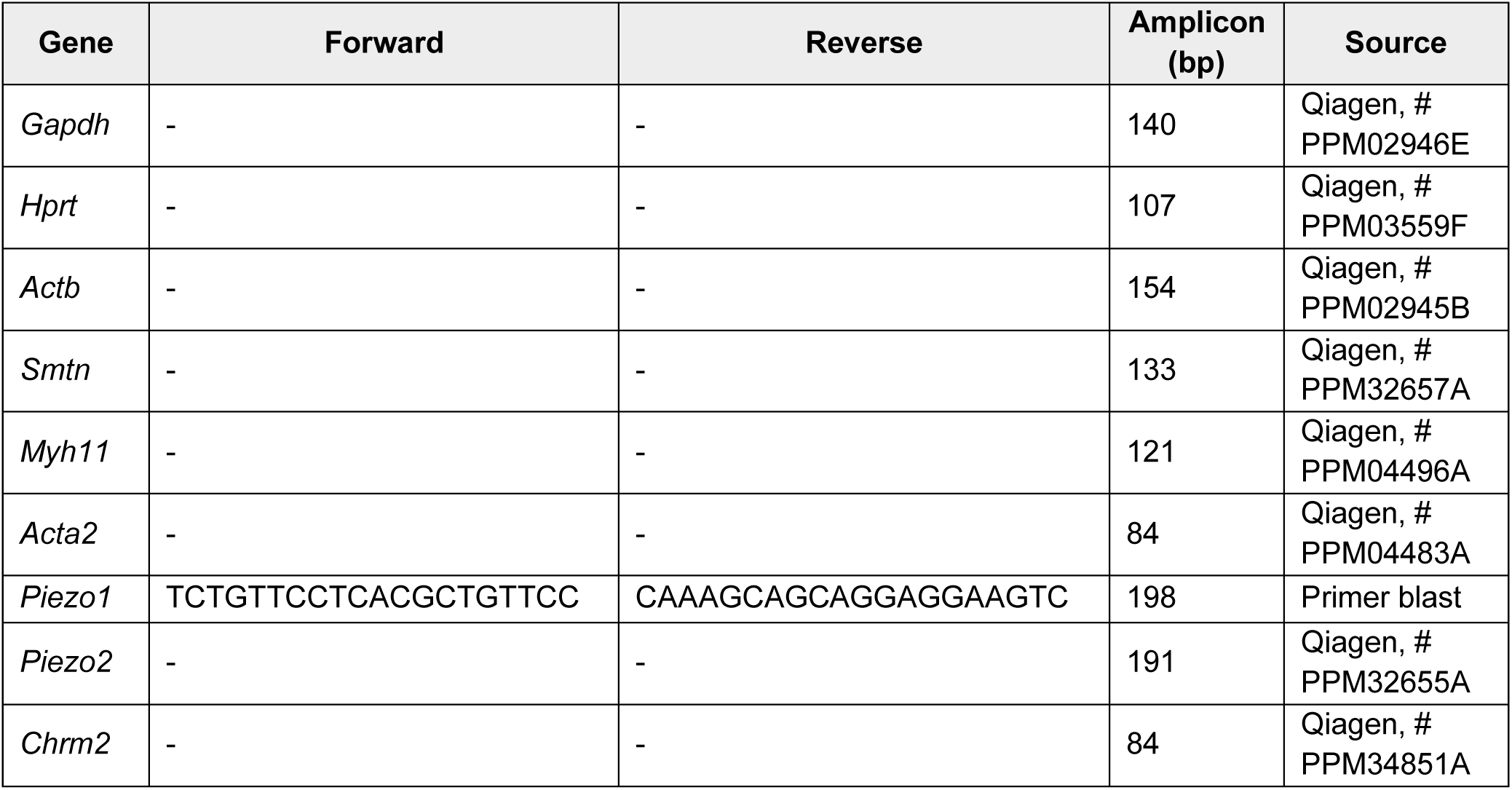
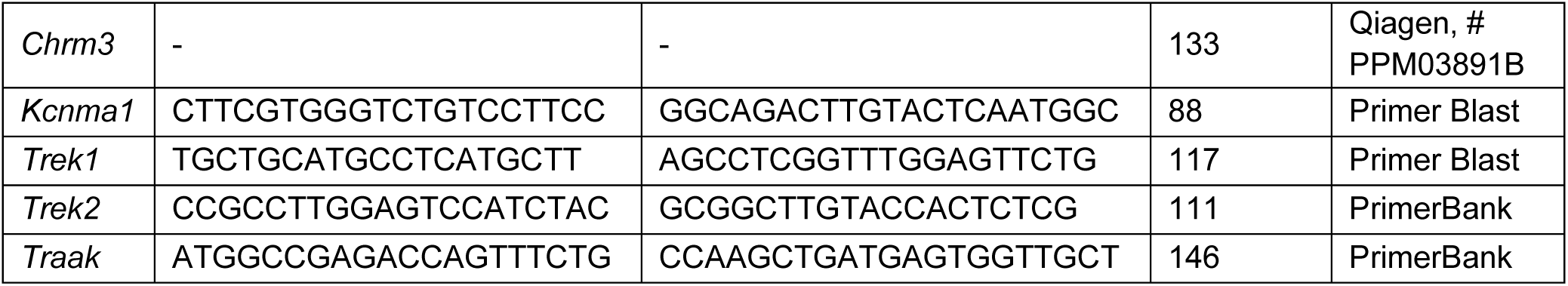
*Mus musculus* RT-qPCR Primers used in this study.

### Liquid chromatography-mass spectrometry

Mouse DRGs dissected from young mice fed a standard chow diet or a margaric acid-enriched diet were rinsed with ice-cold 1X D-PBS (without Calcium or magnesium) and collected in 2 mL screw cap microcentrifuge tubes (Avantor, Cat. #10025 -756). Samples were briefly centrifuged at 800 g for 4 min. and excess supernatant was removed. Samples were then flash frozen in liquid nitrogen. Frozen samples were then shipped to Wayne State University’s Lipidomics core facility where total fatty acids were extracted and quantified. Cellular margaric acid content in each sample was either reported normalized to tissue weight or total protein content, as indicated in each figure. For Figure S4A, animals used were young female mice (∼9-19 weeks).

### Behavioral and Physiological Assays

#### Void spot assay (VSA)

For wildtype C57BL/6J young/aged studies, mice were placed in custom-made bottomless cages that mimic the animals’ home cage environment (11.5” x 7”), set on thick chromatography paper (05-714-5, Fisher Scientific) in a darkened room and allowed to behave freely. Before testing, mice were acclimated to the full testing set-up and behavior room for 2-3 days, 1h each day. On the test day, mice were allowed to behave freely in the test chambers for 4h without access to food or water to avoid accidental water damage to the test papers or biasing where the animals spend time/choose to urinate. Representative animals from each group were placed on a custom-built table with clear acrylic base which is equipped with video cameras at above and below each cage and UV lights placed underneath the cages to allow video recording for movement tracking analysis and urine-spot visualization in real time. At the end of the test, chromatography papers with urine were imaged while illuminated by UV light. Images were thresholded and converted to B&W binary images using ImageJ (v2.16.0/1.54p), and the total number of black pixels was counted. A region of interest corresponding to the center 25% of cage area was drawn and used to quantify the percent of urine signal (pixels) in the center cage arena. For automated void spot count and classification into primary and secondary voids, IMARIS was used (Imaris x64, v11.0.0, Oxford Instruments). Based on evaluation of young, healthy animals’ urinary behavior, the void spot classification threshold was set to 1e-4 µm so that spots >1e-4 µm are considered ‘*primary’*, while spots >1e-4 µm are considered ‘*secondary’*.

For P1KO experiments, VSAs were conducted following best practices as previously described^30,94,95^. Briefly, after a 1 h acclimation period, mice were individually placed in empty cages lined with 3 mm thick chromatography paper (Thermo Fisher Scientific, #057163E). During the 4 h testing period, mice had access to food but not water. Afterwards, chromatography papers were dried and imaged under UV light using a UVP ChemStudio Plus, 815 UV imager (Analytik Jena, Cat. #PN 97-0837-01) equipped with an Auto Chemi Zoom lens (Analytik Jena, #PN 95-0471-01). Images were captured using epi-illumination with UVP Vision Works LS image acquisition software (Analytic Jena, Version #8.20.17096.9551). Void spot analysis was performed in ImageJ using the open-access Void Whizzard software^95^ by an investigator blinded to the treatment conditions. Based on the evaluation of young, healthy animals’ urinary behavior, the void spot spots > 4 cm were considered ‘ *primary’*, while spots > 4 cm were considered ‘*secondary’*.

#### Von Frey

Mice were acclimated to the Von Frey chambers for 30 min. with minimal disturbance, then 10 filaments (0.008 g, 0.020 g, 0.040 g, 0.070 g, 0.16 g, 0.400 g, 0.600 g, 1.000 g, 1.400 g, and 2.000 g) (Aesthesio® Precise Tactile Sensory Evaluator 20 piece Kit, IITC Life Science) were presented to the right hind paw (mid-plantar region), with each filament presented 10 times. A presentation was only counted if the filament exerted enough force on the target area that the filament buckles. Animals’ response/no response to each presentation was recorded, with positive responses including paw shaking, licking, or withdrawal (that is distinct from limb lifting for normal walking).

#### Dynamic touch

Dynamic touch was assessed at baseline (standard chow diet) and after 4-6 weeks on MA-enriched diet. Animals were acclimated to the Von Frey chambers for 30 minutes and 5/0 synthetic brush was gently stroked on the lateral side of the right hind paw in a single motion (heel to tow). Response to each stroke was recorded as follows: 0 = no response, 1 = very short, fast movement/lifting of the paw, 2 = sustained lifting of the paw towards the body for more than 2 seconds, or 3 = flinching, licking or flicking of stroked paw (mechanical allodynia). Scores were reported as the average across 3 trials per mouse (labelled “Response Score”).

#### Gait dynamics analysis

Gait dynamics were quantified using the CatWalkXT (Noldus, v10.6.608.0). First, mice were acclimated to the dark behavior room for 1h with the illuminated surface turned on. To assess each animal’s gait, mice were placed on the illuminated CatWalk walkway and allowed to move freely in both directions and the system automatically video recorded and analyzed the animal’s behavior.

Each test lasted roughly 10 min. or less. Each time the animal crossed the entire length of the walkway was counted as a run. Three compliant runs were analyzed per mouse, where compliance was determined based on the run duration (minimum 0.5s, maximum 5s) and speed variation (maximum 60%). After acquisition, paw positions were manually verified.

#### Gut function

For gut function analysis, mice were acclimated to the behavior room in their home cages for 1h with access to food and water. For testing, mice were them moved to individual, custom-made bottomless cages that mimic the animals’ home cage environment (∼11.5” x7”), set on thick chromatography paper (05-714-5, Fisher Scientific) and allowed to behave freely for 1h. During the 1h test, the number of fecal pellets dropped was recorded, and fecal pellets were collected into air-tight Eppendorf® tubes as soon as they dropped to protect from water loss. The wet weight of fecal pellets was recorded, then the tube with fecal pellets were dried overnight at 100°C in a thermal block/mixer (13687717, Thermo Scientific), with the tube top open to allow water evaporation. Dry pellets weight was recorded the next morning and the change in weight from wet to dry was used to calculate the percent of water content in fecal pellets.

## Electrophysiology

### Cell culture

Primary cultures of mouse DRG neurons were obtained from 8-week-old male C57BL/6 mice (animal numbers/group: 5x Control, 5x MA Diet, 4x MA Media). Mice were anesthetized with isoflurane and then sacrificed by cervical dislocation. DRGs were dissected and kept on ice in Hank’s balanced salt solution 1X (HBSS without CaCl_2_ and MgCl_2_) (Gibco Cat. 14170112). Then DRGs were incubated in 1 mg/mL collagenase B (Roche, Cat. 11088807001) in HBSS, at 37 °C and 5 % CO_2_. After one hour, DRGs were dissociated in medium without serum. The cell suspension solution was centrifuged for 8 min at 800 rpm. The obtained pellet was resuspended in DMEM complete media (Invitrogen, Cat. 11965-118) containing 1% penicillin-streptomycin, 1% MEM vitamin solution (Gibco, Cat. 11380-037), 1% L-glutamine (Invitrogen, Cat. 25030-081), and 10% horse serum (Gibco, Cat. 16050-122). Cells were cultured on glass coverslips pre-treated with poly-L-lysine (Sigma-Aldrich, Cat. P4707). All cultured neurons were used after 18-24 h.

Prior to electrophysiological measurements, DRG neurons were supplemented overnight (≈18 h) with margaric acid, unless otherwise stated. Margaric acid was obtained from (MilliporeSigma, Cat. # H3500). The cultured cells were maintained at 37 °C, 95% relative humidity, and 5% CO _2_.

### Whole-cell patch-clamp & current-clamp

For whole-cell recordings, the bath solution contained 140 mM NaCl, 6 mM KCl, 2 mM CaCl_2_, 1 mM MgCl_2_, 10 mM glucose, and 10 mM HEPES (pH 7.4; 300 mOsm). The pipette solution for voltage-clamp recordings of mechanocurrents contained 140 mM CsCl, 5 mM EGTA, 1 mM CaCl _2_, 1 mM MgCl_2_, and 10 mM HEPES (pH 7.2); and for current-clamp recordings, 140 mM KCl, 6 mM NaCl, 2 mM CaCl_2_, 1 mM MgCl_2_, 10 mM glucose, and 10 mM HEPES (pH 7.4; 300 mOsm). Pipettes were made from borosilicate glass (Sutter Instruments) and were fire-polished before use until a resistance between 3 and 5 MΩ was reached.

During mechanical stimulation, currents were recorded at a constant voltage (−60 mV, voltage-clamp unless otherwise noted) and voltages were recorded without injecting current (current-clamp). Both variables were sampled at 100 kHz and low-pass filtered at 10 kHz using a MultiClamp 700 B amplifier and Clampex (Molecular Devices, LLC). Leak currents before mechanical stimulations were subtracted offline from the current traces and data were digitally filtered at 2 kHz with ClampFit (Molecular Devices, LLC). Recordings with leak currents > 200 pA, with access resistance >10 MΩ, and cells which giga-seals did not withstand at least 6 consecutive steps of mechanical stimulation were excluded from analyses.

### Mechanical stimulation

For indentation assays, DRG neurons were mechanically stimulated with a heat-polished blunt glass pipette (3–4 µm) driven by a piezo servo controller (Physik Instrumente, Cat. #E625). The blunt pipette was mounted on a micromanipulator at an ∼45° angle and positioned 3–4 µm above from the cells without indenting them. Displacement measurements were obtained with a square-pulse protocol consisting of 1 µm incremental indentation steps, each lasting 200 ms with a 2 -ms ramp in 10-s intervals. The threshold of mechano-activated currents for each experiment was defined as the indentation step that evoked the first current deflection from the baseline. For current clamp experiments, the mechanical threshold was defined as the indentation step that evoked the first action potential. Only cells that did not detach throughout stimulation protocols were included in the analysis. The piezo servo controller was automated using a MultiClamp 700B amplifier through Clampex (Molecular Devices, LLC).

### Quantification

Results were expressed as means ± SD (unless otherwise noted). All boxplots show mean (square), median (bisecting line), bounds of box (75^th^ to 25^th^ percentiles), outlier range with 1.5 coefficient (whiskers), and all data points including maximum and minimum. The time constant of inactivation τ was obtained by fitting a single exponential function, Equation (1), between the peak value of the current and the end of the stimulus:

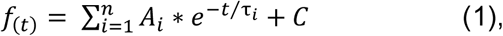

where A = amplitude, τ = time constant, and the constant y-offset *C* for each component *i*. Statistical analyses were performed using GraphPad Instat 3 software. Individual tests are described on each of the figure legends. No technical replicates were included in the analyses.

## Human genomic data

### Sequencing Data Quality Control and Preparation

#### UKB WES Variant and Sample Quality Control

We accessed whole-exome sequencing (WES) data on 470K individuals in the UKB through DNAnexus and applied variant-level quality control (QC) to remove low-quality sites, including genotype quality (GQ) < 20, read depth (DP) < 10, missingness > 5%, and Hardy Weinberg Equilibrium (HWE) p-value ≤ 5e-8. Next, we applied sample-level QC metrics to filter potential problematic samples: 1) genetic ancestry principal component (PCA) analysis, calculated using Somalier v0.2.16^96^, to exclude samples with a European ancestry probability < 0.85; 2) X chromosome heterozygote ratios (xHet), calculated using KING^97^ to exclude sex mismatches, where self-reported males were excluded if xHet exceeded 0.0005 and self-reported females were excluded if xHet was below 0.0005; 3) Somalier relatedness filters to exclude samples with kinship coefficients above 0.2, where a random sample was removed in case-case or control-control relationships and the control sample was removed in case-control relationships; 4) aggregate per sample sequencing statistical outliers, determined by visual inspection of each distribution, including total variants, transition/transversion ratio, singletons, missing calls, heterozygous/homozygous ratios, and indels.

#### AoU WGS Variant and Sample Quality Control

We accessed the v8 AoU srWGS Hail matrix table in June 2025 and extracted exome-restricted variant calls (Gencode v42 basic transcripts ± 15 bp). Next, we removed variants with GQ < 20, call rate < 0.99, allele number < 0.95, and HWE ≤1e-8. We applied sample-level QC using pre-generated auxiliary files and removed samples with low quality (flagged_samples.tsv), European ancestry probability < 85% (ancestry_preds.tsv), related individuals (relatedness_flagged_samples.tsv), where we applied the same case-control removal logic as in the UKB, and aggregate per sample sequencing statistical outliers, determined by visual inspection of each distribution, including total variants, singleton count, heterozygous/homozygous ratio, transition/transversion ratio, and indels.

#### UKB and AoU Variant Annotation

UKB and AoU single-nucleotide variants and indels were annotated with the v94 Ensembl Variant Effect Predictor toolset using GRCh38^98^. We assigned each missense variant a score between 0 and 100 based on the substitution’s predicted impact on overall fitness, using the Evolutionary Action equation^98^. Putative loss-of-function variants, including stop-gained, frameshift, stop-lost, splice-donor, and splice-acceptor, were assigned a score of 100.

### Pathway-Level Enrichment of Ultra-Rare Variants

We applied EA-Pathways^59–61^, a control-free pathway-based association algorithm, to test whether ultra-rare coding variants collectively predispose to neuromuscular bladder dysfunction. For this analysis, ultra-rare variants were defined as population-level doubletons. The empirical null for EA-Pathways is the ultra-rare variant distribution of EA scores observed across the cohort of interest (*e.g.,* bladder dysfunction cases or bladder dysfunction-resistant controls). EA-Pathways is implemented in four steps. First (Step 1), we collapsed ultra-rare variants across each gene and tested their EA score distribution against the cohort background using a Kolmogorov-Smirnov (KS) test. Genes passing Benjamini-Hochberg correction (q < 0.1) were flagged as individually biased and excluded from downstream pathway aggregations. Second (Step 2), the remaining genes were grouped by Gene Ontology (GO) terms^65^ and the EA distribution of the gene set was tested against the null distribution using the KS test in a leave-one-out framework. Genes that increased the functional impact bias of the GO term were designated as “core genes”. Third (Step 3), to further control for inherent functional bias in the GO term gene set, 1,000 simulated pathways ranging between sizes 5 and 50 were generated and optimized as above. GO terms from Step 2 that passed multiple testing correction (Benjamini-Hochberg, *q* < 0.05), contained at least 20 variants, and outperformed the 5^th^ percentile p-value of the same-sized simulated pathways from Step 3 were collected. The final EA-Pathway gene list consisted of all significant genes from Step 1 and the core genes from pathways collected in Step 3.

### Genomic validation analyses

To determine whether carriers of PIEZO variants experience earlier onset of neuromuscular bladder dysfunction, we modeled time-to-event using a Cox proportional hazards framework. Event time was defined as age at first ICD10 diagnosis. Using CoxPHFitter from lifelines v0.27.8, we compared PIEZO-variant carriers with non-carriers, adjusting for sex and ancestry (PC1-3). The resulting plots were visualized using the KaplanMeierFitter function from lifelines. Hazard ratios (HRs) and 95% confidence intervals based on the fitted CoxPH model, adjusted for current age, sex, and ancestry (PC1-3)), are reported.

## QUANTIFICATION AND STATISTICAL ANALYSIS

Statistical analyses were performed using GraphPad Prism (v10.6.1). Data are presented as mean ± SD unless otherwise indicated. The number of independent biological replicates (*n*) and statistical tests used, and significance ranges are provided in the figure legends. Normality was assessed using appropriate tests, and parametric or nonparametric tests were applied accordingly. For comparisons between two groups, paired or unpaired two-tailed Student’s *t* tests or Mann–Whitney tests were used, as appropriate. For comparisons involving more than two groups, two-way ANOVA or two-way mixed-effects models were used, followed by appropriate post hoc tests where applicable. A *p* value < 0.05 was considered statistically significant. Sample sizes were not predetermined using statistical methods but were based on prior studies in the field and practical considerations. Where feasible, investigators were blinded to group allocation during experiments and outcome assessment.

## ACKNOWLEDGEMENTS

The authors thank Drs. Rebeca Caires and Jungsoo Lee for technical support. This work was supported by core facilities including the Wayne State Lipidomics core (NIH grant # S10RR027926 and S10OD032292) and Wisconsin O’Brien Center for Benign Urologic Research (NIDDK grant # U54 DK104310). This work was also supported by the Howard Hughes Medical Institute’s Freeman Hrabowski Scholars Program (K.L.M.) and Cech Fellows Program (T.B.K.), NIH grant numbers R01DK052766 (A.B.), R01DK123549 (A.B.), R00DK128621 (K.L.M.), R25 GM069234 (E.D.L.G.), T32 ES007015 (M.M.R.), The University of Texas Health Science Center at Houston (V.V.), Robert and Arlene Kogod Center on Aging, Mayo Clinic, Career Development Award (V.J.), McNair Medical Foundation (K.L.M.), Pew Charitable Trusts (K.L.M.), and Rita Allen Foundation (K.L.M.)

## AUTHOR CONTRIBUTIONS

Y.M.F.H., V.J., V.V., A.B., and K.L.M. conceptualized the study. Y.M.F.H., V.J., K.W., L.O.R., O.D.S., T.B.K., E.D.L.G., S.P., C.M.C., A.E.M., and M.M.R. performed experiments and analyzed the data. Y.M.F.H., V.J., K.W., and L.O.R. developed the methodology for this work. Y.M.F.H., J.K.A, K.K.S., C.M.V., V.V., O.L., A.B., and K.L.M. supervised the work. Y.M.F.H. visualized and validated data. Y.M.F.H., V.J., K.W., A.B. and K.L.M. wrote the original manuscript draft. Y.M.F.H., V.J., L.O.R., E.D.L.G., J.K.A, K.K.S., V.V., O.L., A.B., and K.L.M. provided edits on the manuscript draft. All authors reviewed the manuscript.

